# REM Sleep Iota Oscillations Across the Lifespan

**DOI:** 10.64898/2026.10.14.759205

**Authors:** Sophia Snipes, Roy Cox, Elena Krugliakova, Salome Kurth, Sven Leach, Andjela Markovic, Valeria Jaramillo, Carina Volk, Fiona Pugin, Ines Wilhelm, Leila Tarokh, Eus van Someren, Reto Huber, Thomas Andrillon

## Abstract

1.

Iota oscillations are a newly discovered marker of wakefulness and rapid eye movement (REM) sleep. So far, iota in REM sleep has only been reported in the surface electroencephalography (EEG) of a small sample of young adults and in intracranial recordings of epileptic patients. Here, we investigated the presence of these oscillations in a large dataset (N=937) of high-density sleep EEG recordings from individuals 7 to 70 years old. We found that iota spectral peaks during REM sleep were between 25 and 40 Hz, slightly faster than in wake (25-35 Hz). Iota power was significantly associated with REM sleep architecture, increasing with both REM duration and across progressive sleep cycles. Iota peaks were observed in individuals of all ages, although the prevalence decreased gradually with age, from 88% in children to 44% in older adults. Most iota spectral peak properties (frequency, amplitude, timing, topography) likewise shifted with age. The most striking effect, however, was in bandwidth, such that children had narrow iota peaks, older adults had broad iota peaks, and 20-30 year olds could have either. It remains unclear whether narrowband and broadband iota are functionally equivalent. On shorter time scales (< 1 year), iota properties remained stable across measurements. Analyses of a separate dataset of monozygotic and dizygotic twins revealed high heritability of all iota peak properties except bandwidth. Overall, these results establish iota as a genetically determined trait-like marker of REM sleep with unique age-related changes. Iota oscillations open new avenues to investigate the interaction between sleep, genetics, development, and aging.

**Teaser:** A new brain wave in REM sleep undergoes a unique transformation in people’s 20’s, possibly reflecting late-stage adult development.

## 2 INTRODUCTION

At first glance, the surface EEG during REM sleep is relatively unremarkable. It is formally described by the American Academy of Sleep Medicine (AASM) as having “low amplitude mixed frequency activity” with occasional sawtooth waves (Berry et al., 2012). However, in practice it is most easily distinguished from other sleep stages by peripheral physiological signals, especially eye movements and reduced muscle tone, and the *lack* of other characteristic EEG events. By contrast, wakefulness is typically dominated by long trains of alpha oscillations (8-12 Hz), and non-REM (NREM) sleep by large slow waves and sporadic spindles (12-17 Hz, i.e. “sigma”). These signals have been extensively investigated, and inform our current understanding of neuronal processes during these states. The absence of such clearly identifiable neuronal features for REM sleep has hampered comparable neurobiological research in this state.

Recently, a new class of oscillations was discovered between 25 and 35 Hz, referred to as “iota” (Snipes, 2025). These oscillations are present during both wakefulness and REM sleep but absent from NREM sleep (Snipes, 2025; Chowdhury et al., 2026). They occur in bursts, more often in proximity to eye movements, and are therefore characteristic of the substage known as phasic REM sleep (Simor et al., 2020). Unlike previously described vigilance markers, iota oscillations have extremely low amplitudes, usually too small for visual detection from the surface. Using EEG, they are most easily identified as peaks in spectral power, but such peaks could only be detected during REM sleep in 7 out of 19 young adults (Snipes, 2025).

These oscillations have been measured more reliably intracranially from the central thalamus of 14 out of 17 adult epileptic patients (Chowdhury et al., 2026), although considering a broader frequency range of 19-45 Hz. The higher prevalence of iota oscillations in intracranial recordings suggests that the lower prevalence in the surface EEG was due to an attenuation of the signal reaching the scalp rather than an absence of iota activity at the source.

In addition to iota oscillations being extremely small, they are at frequencies plagued with overlapping signals and artefacts. To quantify iota spectral activity specifically, it is essential to remove the EEG’s aperiodic background signal (Donoghue et al., 2020; Suppl. Fig. 1.1A), which is highly state dependent (Colombo et al., 2019; Lendner et al., 2020; Schneider et al., 2022; Höhn et al., 2024; Ameen et al., 2025; Rosenblum et al., 2025) and can otherwise mask iota activity when relying on standard spectral bands (Ferri et al., 2000, 2001). Artefacts such as harmonics of lower frequencies, eye movements, heartbeat and muscle tone must also be accounted for. This is done best with high-density EEG, whose greater spatial resolution allows the dissociation between cortical and peripheral signals, and multiple channels provide redundant sources of an otherwise weak and localized signal.

Therefore, to investigate iota oscillations in REM sleep, we pooled high-density sleep EEG data from multiple studies of the University Children’s Hospital of Zurich, and the Netherlands Institute for Neuroscience in Amsterdam, resulting in 937 whole-night sleep recordings from 578 individuals spanning 7 to 70 years of age. These datasets consisted of a combination of healthy controls, as well as patients with attention deficit hyperactivity disorder (ADHD) and insomnia (Suppl. Figure 1.1B).

Our primary aims were to: 1) validate the presence of iota oscillations during REM sleep in surface EEG, using sufficient sample sizes; 2) determine whether variability in iota activity reflected night-to-night differences or could be explained by age, sex, or clinical diagnosis. We further investigated iota in a separate dataset of 31 twin pairs (Markovic et al., 2018) to: 3) evaluate whether iota properties were hereditary.

Oscillatory activity was primarily quantified as individual peaks in periodic spectral power (Suppl. Figure 1.1A). Each peak was defined by its frequency, its amplitude, and its bandwidth. The amplitude of periodic power (units expressed for convenience as “pp”, periodic power, see Methods section 5.2.2) largely reflects the density of a given oscillation in the signal (Snipes et al., 2026), and the bandwidth reflects how precisely that oscillation occurs at the exact peak frequency (resulting in a narrow bandwidth) or not (broad).

## 3 RESULTS

### 3.1 Iota band limits

First, the band limits of iota activity during REM sleep needed to be established without prior assumptions. Spectral power was averaged across channels and epochs, split by sleep stage. The aperiodic signal was removed, leaving periodic power. From this signal, all peaks in each spectrum were detected. To map all the peak properties together, scatterplots in Figure 1 plot peak bandwidths as a function of peak frequency, with point size reflecting peak amplitude. As in Snipes (2025) for wake, a spontaneous cluster emerged in the iota range in REM sleep (Figure 1, left), crucially distinct from the neighboring beta (16-25 Hz) cluster. Fitting a gaussian to the distribution of peaks around 30 Hz, we found that 95% of peaks lie between 25 and 39 Hz (see methods 5.3.1). In wake, the peak was less defined but still present, and the limits were lower, between 26 and 34 Hz, replicating the previous range identified in resting wake EEG (Snipes, 2025). Given that REM sleep iota occupies a broader range, we extended the iota band to between 25 and 40 Hz for subsequent analyses.

**Figure 1:**
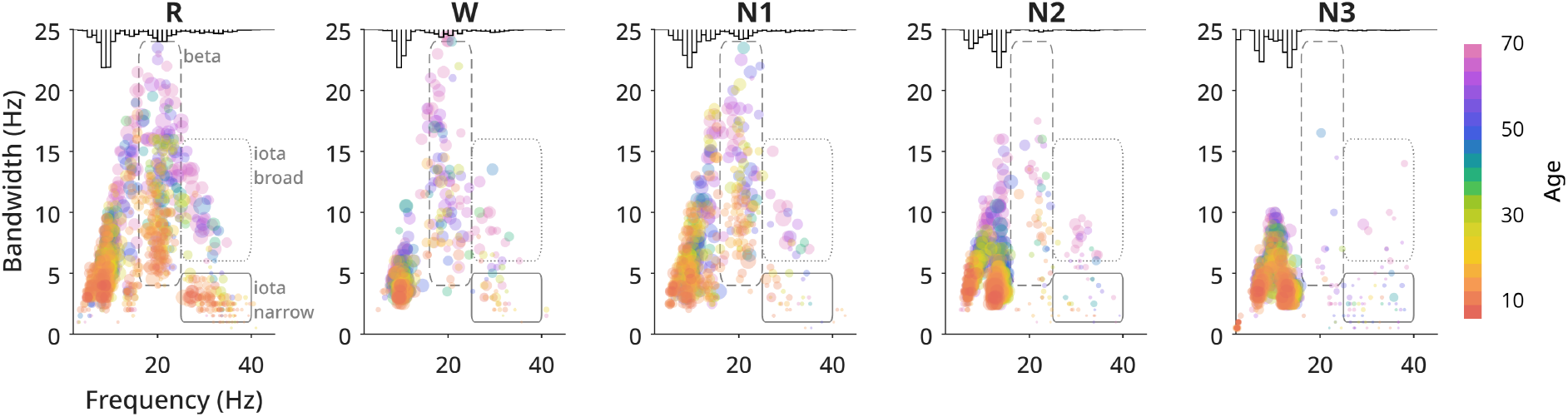
All individuals’ periodic peaks, split by sleep stage. The method for detecting peaks is illustrated in Suppl. Figure 1.1A. Here, the x-axis is the peak frequency, the y-axis is the peak’s bandwidth, the size of each dot reflects the peaks’ amplitude, and the color is the age of the participant. At the top of each plot is a histogram of the total number of peaks per frequency. Peaks were excluded as harmonics if their frequency was exactly two or three times that of a stronger peak at a lower frequency. For participant age distribution, see Suppl. Figure 1.1B. For the same figure including harmonics, see Suppl. Figure 1.2A.

The iota cluster was of narrow bandwidth (1-5 Hz), replicating previous findings (Snipes, 2025), but a second cluster appeared in the same frequency range, with bandwidths between 6 and 15 Hz. This cluster was less distinct from neighboring beta, and was predominantly occupied by adults whereas the narrowband cluster originated from children. The distinction of two clusters by bandwidth is addressed below in Results section 3.4.

Figure 1 excluded peaks which could be harmonics (when a lower-frequency, higher-amplitude peak occurred at a frequency that was one third or one half its frequency). In NREM sleep, without excluding such peaks, there is a cluster of peaks between 20-30 Hz, overlapping with the iota band (Suppl. Figure 1.2A). However, these peaks were tightly related to the frequencies of sigma peaks within each individual, implying they were most often harmonics (Suppl. Figure 1.2B). In addition to these potential harmonic peaks, there remain a few sparse iota peaks in NREM sleep, but they were rare and generally of low amplitude (Figure 1, right; Suppl. Figure 1.2D, Suppl. Figure 2.2). These were most often in NREM2, among older adults, and their peak frequency was unrelated to that of REM iota, unlike the iota peaks in NREM1 (Suppl. Figure 1.2E).

### 3.2 Iota stability across measurements

Using channel-averaged data to detect oscillatory activity as in Figure 1 can be robust to noise but may also miss many low-amplitude signals. For this reason, we developed a more sensitive approach to detect iota peaks, outlined below. As illustrated in the examples of Figure 2A and Suppl. Figure 2.1A-C, sleep spectra across two nights in the same individual are highly preserved, also for iota. We could therefore evaluate the reliability of our new detection method by comparing the stability of iota peak properties across multiple nights within subjects.

**Figure 2:**
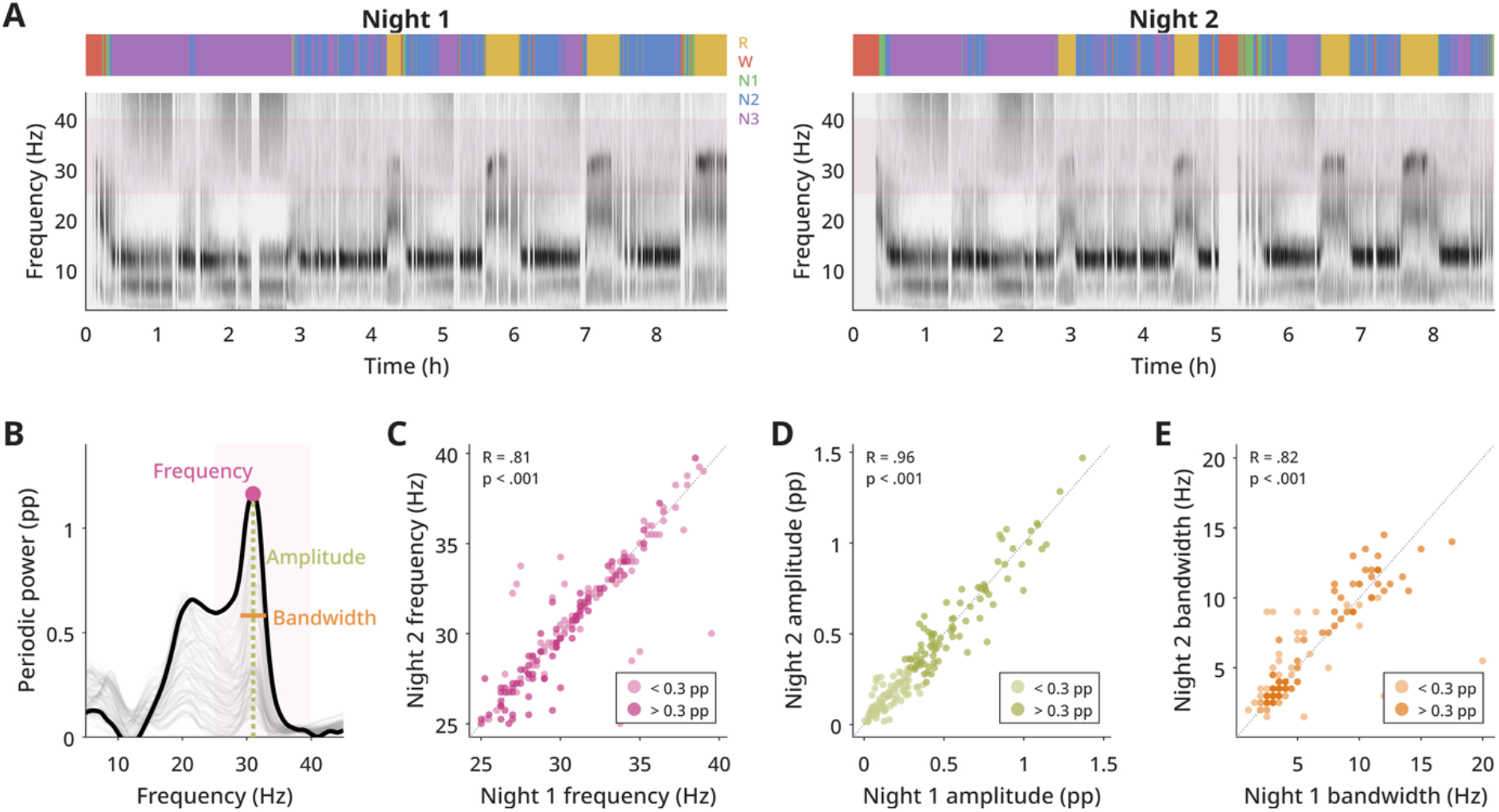
Night to night consistency of iota peaks. A) Time-frequency plots of periodic power from two nights of an example participant. Each night contains a multi-colored bar plotting AASM manual sleep staging. Below it is the spectrogram of periodic power for each epoch, channel-averaged, with darker shades indicating higher amplitudes. A pink band highlights the iota range. Other examples of less clear iota still preserved across nights are provided in Suppl. Figure 2.1. B) The method for detecting iota peaks across channels, using the REM epochs from night 1 of A. Spectral power was calculated on 20 s or 30 s epochs (depending on the original scoring of the dataset), which were averaged, smoothed, and then had the aperiodic component removed with specparam (thin gray lines are periodic power for each channel). Peaks were detected from periodic power (pp) within each standard band (pink patch for iota), and the highest amplitude peak among all channels was selected as the main peak for that sleep stage of that recording (maximum peak is pink circle, thick black line is corresponding channel’s spectra). This peak is defined by its frequency, amplitude and bandwidth. Note, this participant had an exceptionally high-amplitude iota peak (see D). Examples of iota peaks detected in NREM with this method are provided in Suppl. Figure 2.2. C) Comparison in peak frequency of two sessions, color coded by average peak amplitude. Dotted line is the line of equality. Pearson’s correlations and p-values are provided. D) Same as C for peak amplitude. E) Same as C for peak bandwidth.

First, periodic power was calculated separately for each channel, after averaging epochs within each sleep stage. Then, the highest amplitude periodic peak of all channels was selected within the 25-40 Hz range (Figure 2B). A candidate peak was considered a true signal if at least four other channels had peaks within 0.5 Hz of its frequency, each with at least half its amplitude. Iota peaks detected with this approach had the same peak frequency as detected with channel averages (Suppl. Figure 1.2F) but were systematically larger and peaks were detected more often (Suppl. Figure 1.2G). With this method, an iota peak was detected in 62% of all recordings (Suppl. Figure 2.1G).

The absence of a peak in many recordings was most likely driven by the weakness of the iota signal. While the peak amplitudes of wake alpha (mean ± SD: 1.39 ± 0.33 pp) and N2 sigma (0.91 ± 0.18 pp) were approximately normally distributed, REM iota peaks resulted in a half-normal distribution with a floor at 0 (0.34 ± 0.29 pp; Suppl. Figure 2.1D). This underscores that iota activity is often at signal-detection threshold levels, and therefore for many individuals it is likely drowned out by the overall background EEG signal, even in otherwise clean data.

Nevertheless, even the smallest iota peaks remained consistent night to night in frequency (Figure 2C), amplitude (Figure 2D) and bandwidth (Figure 2E). Intraclass correlation coefficients (ICC) for iota frequency remained around .9 (considered excellent (Koo and Li, 2016)) for peaks above 0.3 pp, and only dropped to ∼.8 (still considered good) when including amplitudes below 0.3 pp (Suppl. Figure 2.1E). 92% of participants had iota peak frequencies within 1.5 Hz across two nights when mean amplitude was > 0.3 pp, and this dropped to 85% for peaks < 0.3 pp (Suppl. Figure 2.1F). Only 15% (49/313) of individuals had a peak in just one session, and this peak had an average amplitude of 0.14 ± 0.16 pp. Thus, iota peaks were a highly stable trait across nights, and only individuals with the smallest peaks had less consistency or could sometimes miss having a peak detected in a second night.

### 3.3 Iota specificity to REM sleep

Having established a reliable definition of iota, we needed to confirm that iota activity truly characterized REM sleep in contrast with NREM sleep, and was independent from general EEG activity. Figure 3 compares spectral activity across sleep stages, divided by bands, starting with an illustrative example (Figure 3A-B), and then averaging all recordings (Figure 3C; all pairwise statistics for log power, periodic power, and peak amplitudes in Suppl. Tables 1-3).

**Figure 3:**
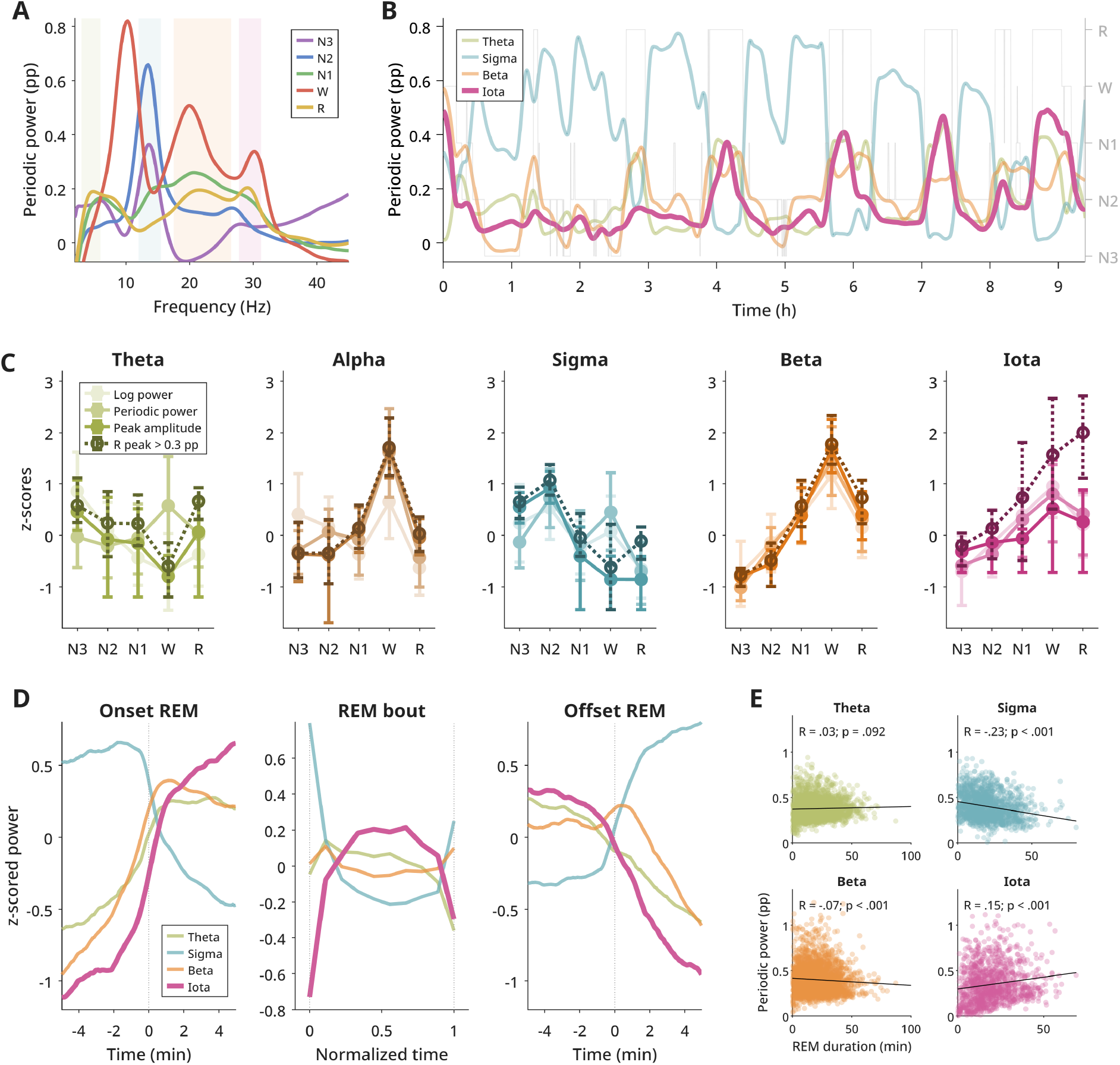
Specificity of iota to REM sleep. A) Channel-averaged periodic power for each sleep stage from an example recording. Colored patches reflect the custom bands used in B. B) Smoothed timecourse of periodic power from the example recording in A, averaged into custom bands (individual peak frequency ± peak bandwidth/2) from the top 10 channels of each spectral peak. Gray lines provide AASM sleep staging. C) Average power of all electrodes for each sleep stage, split by band. Each plot provides average log power and periodic power using canonical bands (theta, 4-8 Hz; alpha, 8-12 Hz; sigma 12-16 Hz; beta 16-25 Hz; iota 25-40 Hz), compared with periodic peak amplitudes, and periodic peak amplitudes selecting only recordings with REM peak > 0.3 pp. Data was z-scored across observations for visualization purposes (not for statistics). For peak amplitudes, if no peak was detected, the value was set to 0. Error bars depict interquartile ranges. D) Average smoothed (1 min moving mean) periodic power at transitions into REM sleep (left) and out of REM sleep (right), z-scored, using custom bands. Only participants with peaks > 0.3 pp for a given band were included. Middle panel is z-scored power during REM bouts of all lengths > 5 min, normalizing time into 10 quantiles. Note, the data used in the figure includes all participants, whereas the statistical text provided in the text includes only participants with both iota and beta peaks. E) Correlation between REM duration and REM average periodic power, per sleep cycle, pooling all cycles of all recordings for which the overall night’s REM peak of a given band was > 0.3 pp. Pearson’s correlations and p-values are provided in each figure. Note, this correlation does not take into account repeated measures, and is intended only as a rough comparison between bands; repeated measures are instead addressed in Results section 3.4.1.

Iota peak amplitudes were highest in wake, but closely followed by REM sleep and then progressively lower with increasing NREM sleep stage, the difference between REM and NREM being statistically significant (R vs W peak amplitudes: t = −1.23, p = .279, d = −.08, df = 116; R vs N1: t = 7.37, p < .001, d = .43, df = 143; R vs N2: t = 13.53, p < .001, d = .87, df = 274; R vs N3: t = 14.98, p < .001, d = 1.18, df = 251). Beta activity was similar, although the difference between wake and REM sleep was also statistically significant (R vs W: t = −24.16, p < .001, d = −1.62, df = 232). As expected, alpha activity was strongest in wake, and sigma activity in NREM sleep (Suppl. Table 3). Theta periodic peak amplitudes were actually highest during N3 (R vs N3: t = −6.14, p < .001, d = −0.38, N = 360), in contrast to what is observed in rodents, where theta is a defining marker of REM sleep (Adamantidis et al., 2019). Overall, beta and iota were the most characteristic oscillations of REM sleep, while both were also found in wake.

Given that beta and iota are adjacent bands, to consider them functionally distinct they must exhibit sufficiently distinct behavior. This can be seen already in the example Figure 3B, where the traces of iota and beta fluctuate independently, even though both peak around REM sleep. Figure 3D provides the group average transitions of periodic power into and out of REM sleep. Beta activity anticipates the rise of iota by 1.25 min (t = 15.13, p < .001, d = 1.11, N = 286), and decreases after iota by 1.08 min (t = 10.04, p < .001, d = .81, N = 210).

Beta power peaks shortly after the transition into REM sleep, whereas iota continues to rise across the first five minutes of REM sleep (Figure 3D, left). The middle panel of Figure 3D illustrates how iota activity peaks in the middle of the REM bout (defined as at least 5 min of continuous REM sleep), and decreases before bout offset. This suggests that the maximum iota power depends on REM duration, which is confirmed in Figure 3E, where REM iota power was found to be significantly correlated with REM duration within each sleep cycle, unlike other bands. Conversely, beta activity seems to preferentially occur around the transitions themselves and is (slightly) negatively correlated with REM duration.

#### 3.3.1 Wake iota vs REM iota

Wake EEG recorded in bed around sleep is not as reliable as periods of sleep themselves; in some individuals it can be extremely brief, fragmented, and is often plagued by movement and muscle artefacts. Nevertheless, we took the opportunity to compare iota peak frequencies in wake and in REM sleep in the 154 individuals which had sufficient wake EEG, sufficient REM EEG, and a REM iota peak.

Wake and REM iota peak frequencies were significantly but not substantially correlated with each other (R = .30, p = .001, N=117). REM iota peak frequencies were not correlated with either wake alpha (R = .05, p = .526, N = 150) nor N2 sigma (R = −.09, p = .108, N = 346), ruling out that the correlation with wake iota was simply due to generally “faster” or “slower” brains. The REM iota peak was usually faster than in wake (mean diff = 1.09 Hz; t_paired = 3.29, p = .001, d = .36, df = 116). As noted earlier, the range of peak frequencies is lower in wake than in REM sleep (35 Hz vs 40 Hz max). For participants with a fast REM iota peak (outside of the wake range; ≥ 35 Hz), 45% (10/22) did not have a wake peak at all. By comparison, only 20% (27/132) of participants with a slower REM iota peak (inside the wake range; < 35 Hz) did not have a wake peak (chi2 = 6.46, p = .011, OR = 3.24). The mean wake peak frequency was 29.6 ± 2.8 Hz for participants with high REM peaks, well below the lower bound of 35 Hz. Furthermore, the mean REM peak amplitude for those high-frequency iota peaks was 0.13 ± 0.17 pp. Altogether, this indicates that REM sleep iota peaks are typically faster than in wakefulness, but additionally, individuals with high-frequency REM peaks are less likely to have a wake iota peak at all, possibly due to their lower amplitudes not making it above the background signal during wakefulness.

### 3.4 Age effects on REM sleep iota

All spectral properties changed with age (Suppl. Figures 4.1-4.9), including iota in REM sleep (Figure 4). The prevalence of detected iota peaks decreased with age, although curiously the prevalence of high amplitude peaks increased (Figure 4A). While a peak was found in 88% of children under 12 y.o., this dropped to 44% in adults over 55 y.o.

**Figure 4:**
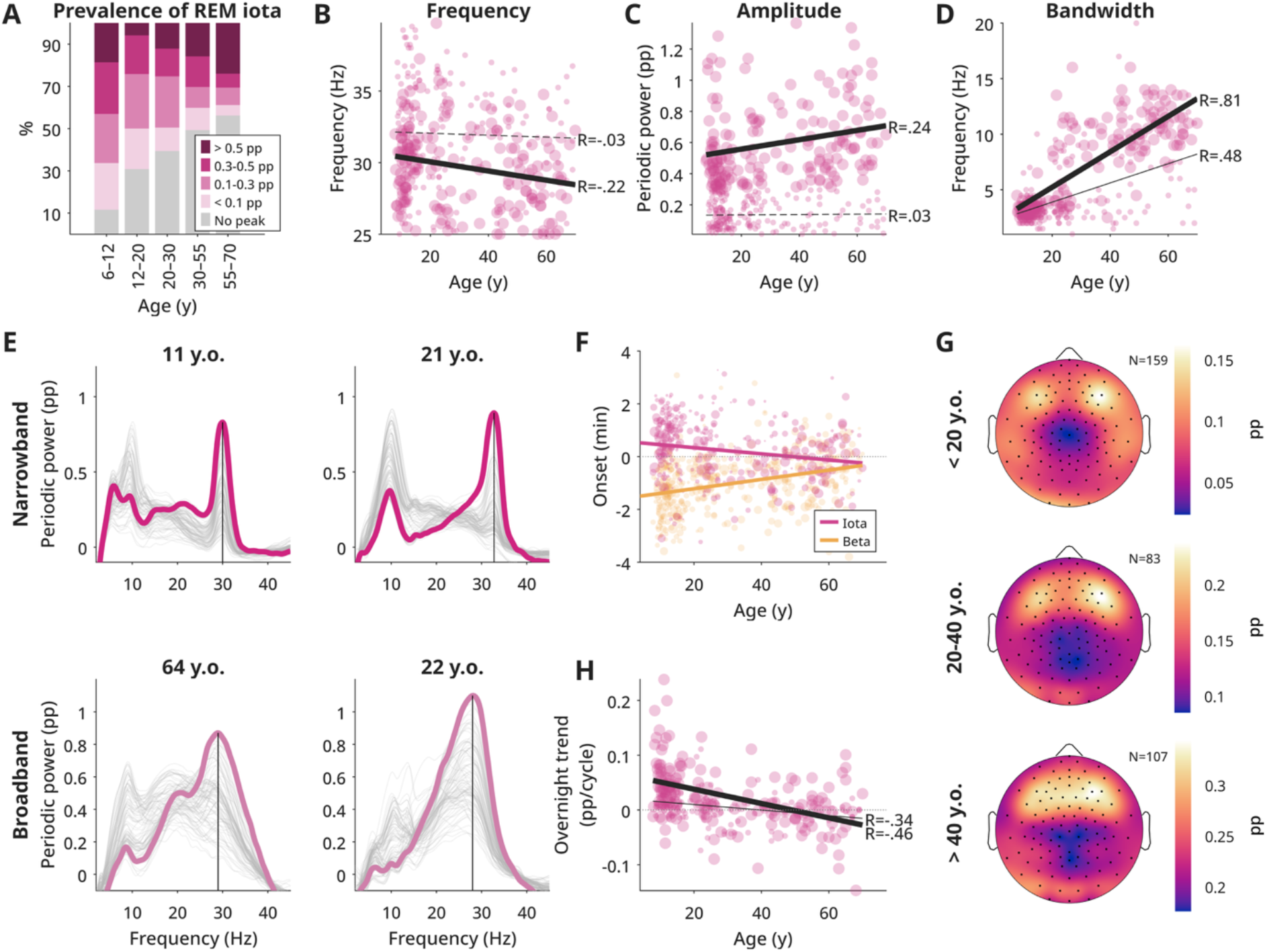
Age effects of iota. A) Proportion of individuals with iota peaks of different amplitudes (or no peak), split by age. Suppl. Figure 4.1 provides the same for all sleep stages and bands. B) Correlation between age and iota peak frequency, C) amplitude and D) bandwidth. Dots are large if peak amplitude was > 0.3 pp, and small if < 0.3 pp. The thick regression line is for large amplitudes, the thin line for small. The line is dashed if the Pearson’s correlation is not statistically significant. Suppl. Figures 4.2-4.4 provide the same for all sleep states and bands. E) REM sleep periodic power in four examples of different ages, two narrowband, two broadband (y.o. = years old). F) Midpoint in time of iota and beta band transition into REM sleep as a function of age. Dots are scaled to peak amplitudes. G) Topographies of custom iota periodic power, split by age. The number of participants in each average is provided in the top right. All participants with an iota peak in REM sleep were included. Suppl. Figures 4.5-4.9 provide the same for all bands and stages. Given that both bandwidth and dataset changes with age, Suppl. Figure 4.10 provides the average topographies of narrowband and broadband iota power split by dataset for 20-40 y.o. This indicated that the more ovaloid peaks in >20 y.o. are due to the different EEG nets. H) Change in periodic iota power across sleep cycles, plotted against age. Linear regression coefficients (pp/cycle; positive values indicate increasing power) were calculated by fitting mean REM custom iota power against cycle number per night. Size, color and regression lines are the same as B-D. Only cycles with more than 10 min of REM were analyzed, and only nights with at least 2 valid cycles were included.

Peak iota frequency decreased (Figure 4B), possibly contributing to the decrease in prevalence as slower peaks fell below the 25 Hz lower iota limit. Surprisingly, amplitudes significantly increased (Figure 4C), and it was primarily small iota peaks <0.3 pp that disappeared with age (Figure 4A).

Most notably, peaks increased substantially in bandwidth with age (Figure 4D). While the changes in amplitude and frequency were gradual, the change in bandwidth was abrupt. Children almost exclusively had narrow peaks (bandwidth < 6 Hz): 97% (155/159), rising to 98% (65/66) when considering only peaks with amplitudes > 0.3 pp. Among adults over 30, only 17% (18/107) had narrow peaks, falling to 1% (1/72) when considering only peaks with amplitudes >0.3 pp. Those aged 20-30 showed a bimodal distribution, with 57% (47/83) having narrow peaks, falling to 42% (15/36) for amplitudes >0.3 pp. There was no clear gradient within this young adult group; either young adults had spectra like children, or like older adults (Figure 4E). This jump in bandwidth is specific to iota: almost all bandwidths increased linearly with age (Suppl. Figure 4.4), but these increases were gradual.

Broadband iota becomes less distinguishable from beta. The blurring between beta and iota with age further increases regarding timing into REM sleep (Figure 4F). While in children iota activity followed beta by ∼2 min, in the oldest adults they were nearly simultaneous. This was due to both a gradual anticipation of iota and a delay of beta, so the effect cannot be tied to the abrupt change in iota bandwidth. The effect was mirrored at REM offset (data not shown).

Furthermore, the topography of iota changed slightly with age. In children and young adults, iota appears as two frontal spots, peaking over channel F6. Iota was less localized in older adults, where the two frontal spots became bridged (Figure 4G). Curiously, while in children beta activity showed the same average topography as iota (frontal bilateral), in older adults, beta power was highest over bilateral parietal areas (Suppl. Figure 4.5). Thus, while the beta and iota spectral peaks tend to merge with age, they have different scalp topographies in older adults, reflecting distinct cortical sources.

#### 3.4.1 REM cycle analysis

Figure 3B revealed a steady increase in peak iota power with successive REM sleep cycles. While this could be explained by iota activity increasing with time asleep (or changing with circadian phase), it could also be explained by changes to REM bout duration to which iota is correlated (Figure 3E), and which also increases across the night (Feinberg, 1974). To test this, we used linear mixed effects models to disentangle these effects with the following equation: CustomPower_Iota ∼ CycleIdx * Bandwidth + REMDur * Bandwidth + Age + Lab + (1|Participant) + (1|Participant:Recording). CustomPower_Iota was the average iota power in the custom band around the detected peak frequency. REMDur was the amount of REM minutes in a given cycle (CycleIdx), both of which were z-scored. Bandwidth was here a binary category of narrowband (iota peak bandwidth ≤ 6 Hz) or broadband (iota bandwidth > 6 Hz). The full model output is provided as extended data.

We found significant main effects of bandwidth (broad referenced to narrow: β = 0.0862, t = 6.86, p < .001, df = 2432), sleep cycle (β = 0.0327, t = 18.26, p < .001, df = 2432) and REM duration (β = 0.0199, t = 10.90, p < .001, df = 2432), as well as a significant negative interaction with bandwidth for sleep cycle (β = −0.0437, t = −11.93, p < .001, df = 2432), but no significant interaction with bandwidth for REM duration (β = −0.00181, t = −0.48, p = .631, df = 2432). This means that iota activity increases across sleep cycles, and independently also increases with REM duration, and the relationship to sleep cycle is diminished or even reversed in broadband iota (Figure 4H). This increase across sleep cycles mirrors an increase in rapid eye movement density across the night (Aserinsky, 1971; Khalsa et al., 2002), further connecting at least narrowband iota and eye movements as complementary markers of phasic REM sleep (Simor et al., 2020; Snipes, 2025; Chowdhury et al., 2026).

#### 3.4.2 Group differences on REM sleep iota activity

While the relationship between age and iota activity was readily apparent, the data further allowed investigations into influences of biological sex, as well as ADHD in children and insomnia in older adults (these were the objects of study of the original datasets). To disentangle these effects, we conducted linear mixed effects models separately for the two datasets (Amsterdam: > 18 y.o., insomnia; Zurich: < 27 y.o., ADHD), with the following formula: PeakMeasure ∼ Age + Sex + Diagnosis + (1 | Participant).

In the Zurich children’s dataset, there was a significant effect of age on the prevalence of iota (β = −0.169, t = −5.28, p < .001, df = 491), iota peak amplitude (β = −0.00930, t = - 2.97, p = .003, df = 362; notably decreasing despite increasing across the whole dataset in Figure 4C), iota peak bandwidth (β = 0.140, t = 5.38, p < .001, df = 362), but not frequency (β = 0.0877, t = 1.81, p = .071, df = 362). In the Amsterdam adult dataset, there was a significant effect of age on the prevalence of iota (β = −0.0215, t = −2.35, p = .019, df = 438), iota peak amplitude (β = 0.00411, t = 2.42, p = .016, df = 214), iota peak bandwidth (β = 0.0828, t = 4.23, p < .001, df = 214), but not frequency (β = −0.0168, t = - 0.88, p = .380, df = 214). The significant negative correlation of frequency with age in Figure 4B only emerges when combining datasets and excluding low-amplitude peaks.

The effect of sex was not significant in the Zurich children’s dataset for any outcome measure (prevalence: β = −0.232, t = −0.75, p = .455, df = 491; frequency: β = −0.00480, t = −0.01, p = .992, df = 362; amplitude: β = −0.0594, t = −1.68, p = .095, df = 362; bandwidth: β = −0.112, t = −0.44, p = .657, df = 362). In the Amsterdam adult dataset, there was a significant effect of sex on iota peak amplitude (female as reference: β = −0.108, t = −2.13, p = .034, df = 214), but not prevalence (β = 0.0391, t = 0.15, p = .883, df = 438), frequency (β = 0.515, t = 0.92, p = .358, df = 214), or bandwidth (β = −0.974, t = −1.70 p = .091, df = 214).

Given that the effect of sex on amplitude was also trending for children, we age-matched male and female participants across both datasets, and performed unpaired t-tests between sexes, confirming a significant but small effect of sex, such that iota amplitudes were larger in females, but the effect was small (male: 0.30 ± 0.27 pp; female: 0.38 ± 0.28 pp; t_unpaired = 2.21, p = .028, d = .29, N = 226).

The effect of ADHD was not significant for any measure (prevalence: β = −0.686, t = −1.76, p = .079, df = 491; frequency: β = −0.532, t = −0.95, p = .341, df = 362; amplitude: β = 0.0294, t = 0.72, p = .475, df = 362; bandwidth: β = −0.0647, t = −0.22, p = .828, df = 362).

The effect of insomnia was not significant for any measure (prevalence: β = 0.220, t = 0.87, p = .387, df = 438; frequency: β = −1.07, t = −1.96, p = .052, df = 214; amplitude: β = 0.0666, t = 1.33, p = .183, df = 214; bandwidth: β = 0.736, t = 1.31, p = .192, df = 214).

### 3.5 Heterogeneity in iota topographies

In addition to spectral peak properties, iota activity differed across individuals in topography. While the group average consisted of bilateral frontal spots (Figure 4G), only around 34% of participants had such an iota topography (Figure 5A, Frontal Bilateral), with an additional 13% having only one of the two front spots (Frontal Unilateral). 16% of individuals had only one central frontal spot (Frontal Midline), although these increased in prevalence with age (Figure 5B), contributing to the older adults’ bridged average topography in Figure 4G. Correspondingly, these recordings were almost exclusively broadband (Figure 5C, right). Similarly, frontal unilateral topographies were more often found in adults, and more often broadband.

**Figure 5:**
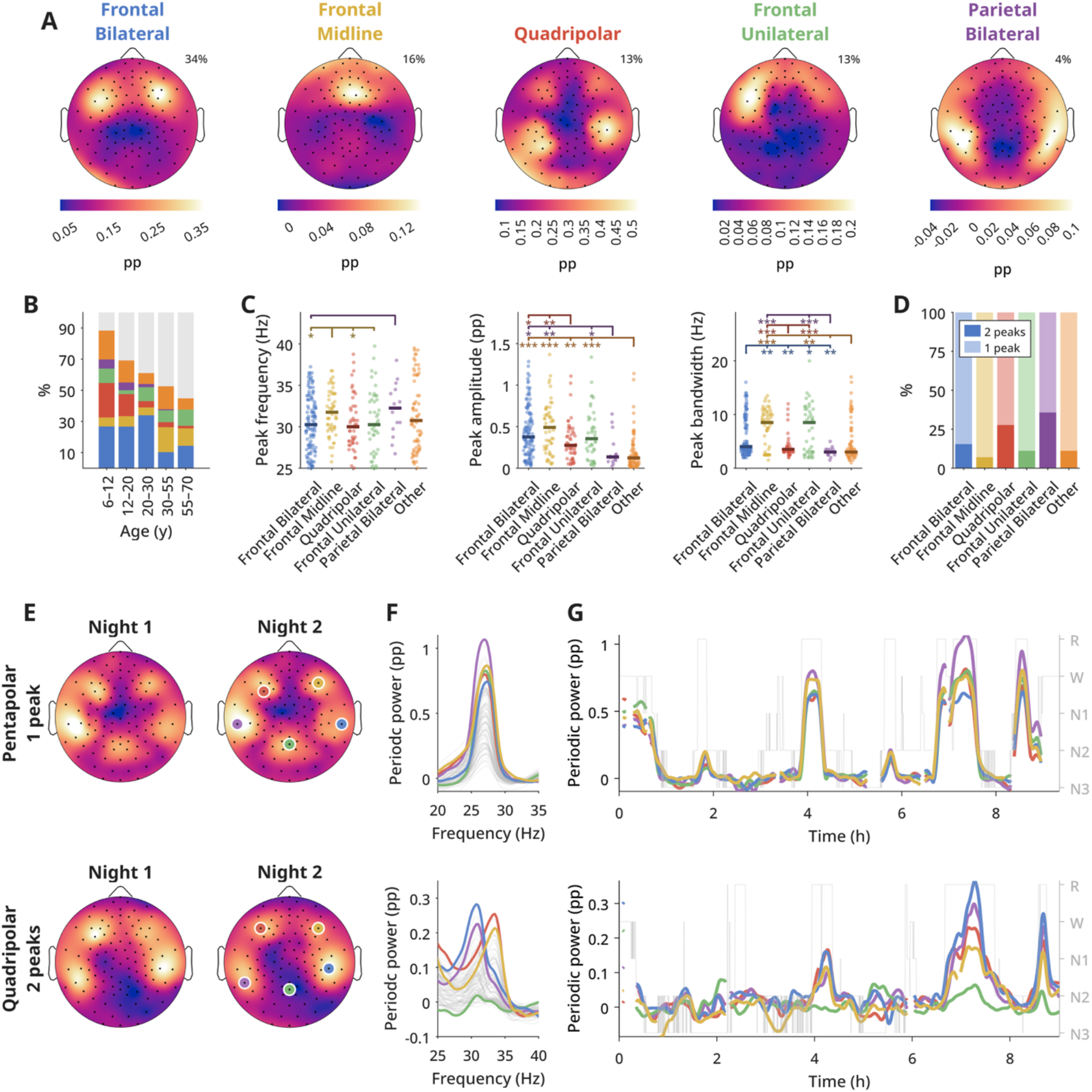
Iota topography variants. A) Individual examples of top 5 topography variants of REM sleep iota periodic power. Percentage of all iota topographies for a given type are provided in the top right corner. B) Distribution of topography variants, split by age. Colors are the same as in C and D: blue = frontal bilateral; yellow = frontal midline; red = quadripolar; green = frontal unilateral; purple = parietal bilateral; orange = other; gray = no iota peak. C) Iota peak properties split by topography variant. Stars indicate statistically significant differences between groups, from unpaired t-tests, FDR corrected (* p < .05, ** p < .01, *** p < .001). D) Proportion of recordings with at least 2 peaks in the iota range per topography type. E) Example of pentapolar (top) and quadripolar (bottom) topographies, across two nights. Colored circles indicate channels highlighted in F & G, selected as the highest amplitude channel of a given pole. F) Periodic spectra of all channels from the examples in E, Night 2. Colored lines are channels highlighted in E. G) Smoothed timecourses of the highlighted channels of the two example participants.

Curiously, 13% of individuals had both two frontal and two parietal spots (Quadripolar). These topographies were primarily found in children and were almost exclusively narrowband. 4% had only the parietal peaks (Parietal Bilateral), which was the average configuration for resting wake (Snipes, 2025). Other configurations covered < 2% of participants, and 20% did not have clearly classifiable topographies (Suppl. Figure 5.1), although most of these had low amplitude peaks (Figure 5C, middle) and thus were easily affected by residual artefacts. Similar variants in topography were also found in wake iota (Snipes, 2025), and NREM sleep spindles (Cox et al., 2017).

Iota topographies were highly consistent across nights. To quantify this, we ran Pearson’s correlations of custom iota periodic power across channels, between two nights of each individual, resulting in an average r of .88 (N = 172; using Fisher’s z transformation as in (Markovic et al., 2018)). This significantly exceeded any similarity between different individuals (r = .30; 10,000 random re-pairings, z_perm = 40.59, p < .001). Figure 5E provides two examples, one of a rare “pentapolar” variant, the other a quadripolar variant, both with nearly identical topographies across measurements. Most night to night differences were due to the relative strength of any given iota “pole”, which could sometimes disappear entirely.

A final variant of iota activity discovered in this dataset was the presence of multiple iota peaks in the spectrum, although rarely more than 2 (Figure 5F, bottom). The likelihood of multiple peaks differed by topography (Figure 5D) and was most common in quadripolar and parietal recordings (and therefore in children). As seen in the example figure, the parietal sources could have different frequencies from the frontal ones (although not always, as in the pentapolar example). Older adults could also present multiple iota peaks from different channels (e.g. in Figure 4E), however the broader bandwidths make such distinctions in peak frequency less clear. Based on visual inspection, iota periodic power from these multiple topographical sources and frequency peaks were modulated largely in synchrony with each other, albeit with different channels reaching different peak amplitudes (Figure 5G).

### 3.6 Iota in twin children

To investigate whether iota properties could be explained by hereditary factors, we analyzed a separate dataset of 58-channel sleep EEG in twin children (9-15 years old), including 18 monozygotic (MZ, i.e. “identical” twins) and 13 dizygotic pairs (DZ, “fraternal” twins). This dataset has previously been analyzed (Markovic et al., 2018, 2022; Rusterholz et al., 2018), and the authors found in fact strong evidence of heritability in these higher frequency ranges, however without yet separating the periodic from aperiodic signal.

Iota peaks were detected in 38/62 (∼61%) individuals (Figure 6A). Iota frequency and amplitude were more highly correlated in MZ compared to DZ twins, but not bandwidths (Figure 6B). Similar results emerged when comparing ICC within MZ vs DZ twins. REM iota peak frequency showed an ICC(1,1) of .99 in MZ twins (95% CI [.94, 1.00]; N = 8 pairs) and .44 in DZ twins (95% CI [-.33, .87]; N = 7 pairs), considered excellent and poor, respectively. Iota peak amplitudes showed an ICC(1,1) of .88 in MZ twins (95% CI [.72, .95]; N = 18 pairs) and .18 in DZ twins (95% CI [-.38, .64]; N = 13 pairs). Bandwidths instead showed an ICC(1,1) of .04 in MZ twins (95% CI [-.61, .68]; N = 8 pairs) and .13 in DZ twins (95% CI [-.60, .76]; N = 7 pairs).

**Figure 6:**
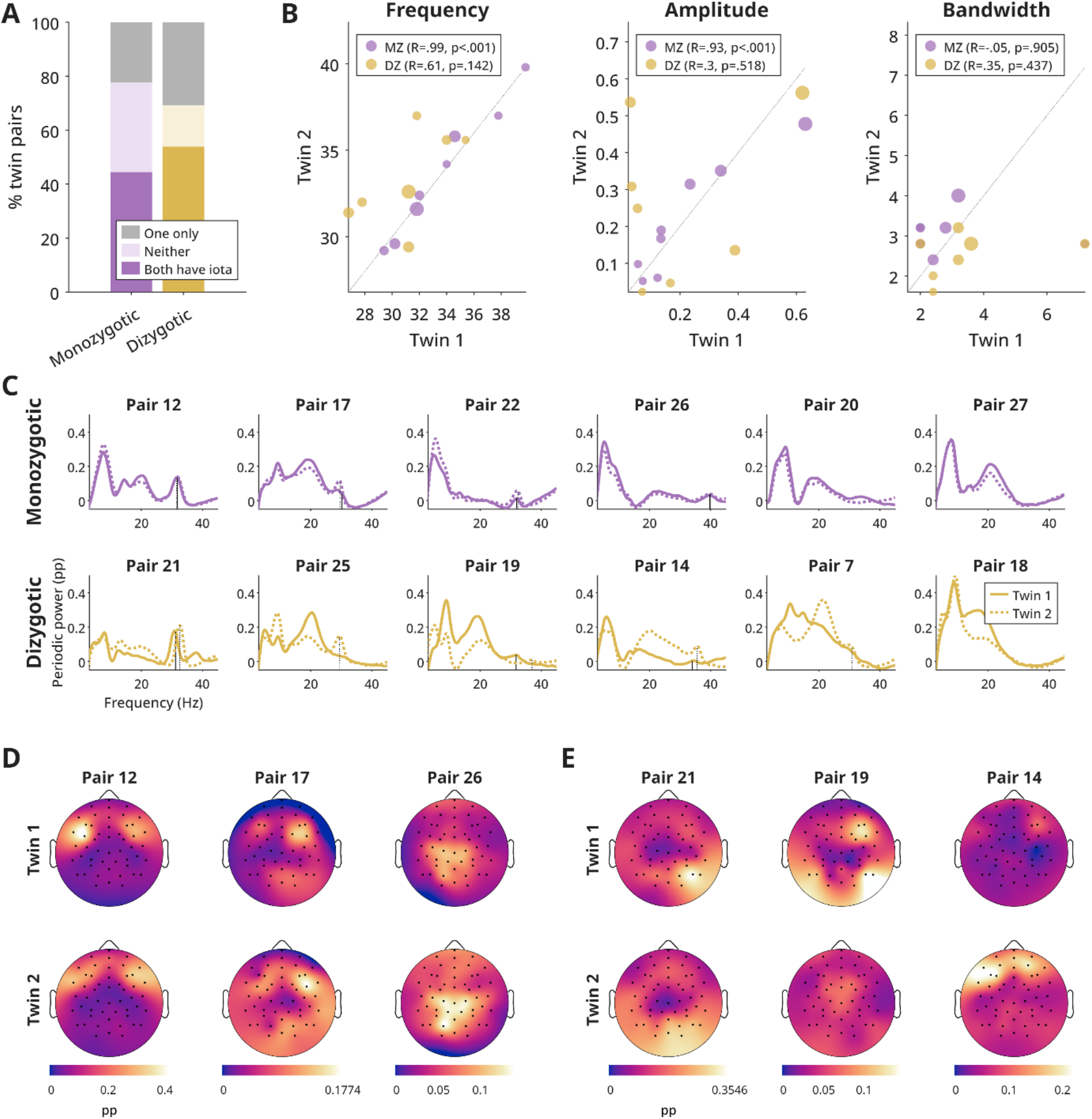
Iota activity in twins. A) Proportion of monozygotic (MZ, identical) and dizygotic (DZ, fraternal) twin pairs in which both have iota peaks (dark color), neither have a peak (light color), or only one twin has a peak (gray). B) Relationship of peak measures between twin pairs. Dot size reflects average peak amplitude. Pearson’s correlations provided for each zygosity. C) Examples of periodic spectra among twin pairs. Iota peaks are marked with a black vertical bar. D-E) Examples of iota topographies (custom frequency band) for monozygotic (D) and dizygotic (E) twin pairs.

We used structural equation modelling (SEM) to determine whether genetic (A), environmental (C) or unique and measurement error (E) components best explained twin covariance of iota peak properties. The ACE model was compared to reduced models (ACE, AE, CE), and the model with lower Akaike Information Criterion (AIC) was selected.

For frequency, the AE model had the lowest AIC (132.18; N_MZ = 8 MZ pairs, N_DZ = 7 pairs), with A = .98 and E = .02. Likewise for amplitudes, the AE model had the lowest AIC (-58.81; N_MZ = 18 pairs; N_DZ = 13 pairs), with A = .92 and E = .08. For bandwidths, the CE model had the lowest AIC (104.57; N_MZ = 8 MZ pairs, N_DZ = 7 pairs), with C = .01 and E = .99. Therefore, heritability is supported for iota frequency and amplitude, but not bandwidth.

Iota topographies also showed stronger similarity within MZ compared to within DZ twins (Figure 6D,E), which is even more striking given the heterogeneity of iota sources. Mean REM iota topographic similarity, assessed with Pearson’s r correlations across channels (see (Markovic et al., 2018)), was for MZ R = .73 ± .13 (mean ± SD; N = 8 pairs), and for DZ R = .40 ± .28 (N = 7 pairs), and R = .19 ± .40 in unrelated participants (N = 615 pairs). Using Welch’s t-test, the difference between MZ and DZ was statistically significant t(11.61) = 2.96, p = .012. Using Falconer’s estimates, heritability (h2) was .67 and environmental influence (c2) was .06, again supporting heritability.

## 4 DISCUSSION

With a new oscillation like iota, the possible research directions are infinite. The goal of this investigation was to narrow down where to start by identifying *who* has iota activity, and how to explain its most apparent heterogeneity. We developed a relatively simple, robust algorithm for low-amplitude spectral peak detection, relying on the redundant information present in high-density EEG (Figure 2B), that could readily be applied to large datasets. Across 578 participants between 7 and 70 years old, this approach identified iota peaks at 25-40 Hz during REM sleep in approximately 60% of recordings. Even the smallest peaks showed consistent frequency, amplitude, and bandwidth across multiple within-subject recordings (Figure 2C-E, Suppl. Figure 2.1), indicating a stable signal.

Having identified iota peaks, we then questioned whether iota activity was truly characteristic of REM sleep and whether it meaningfully differed from other oscillatory activity, especially neighboring beta. We confirmed iota activity was specific to both wakefulness and REM sleep, and mostly absent in NREM sleep (Figure 1). NREM “iota” peaks could sometimes be found, but were most often harmonics (Suppl. Figure 1.2) and significantly of lower amplitude than in REM sleep (Figure 3C, Suppl. Figure 1.2D, Cohen’s d R vs N2 = 0.87). Beta activity was also specific to wakefulness and REM sleep, but the dynamics differed from iota; beta activity began increasing minutes before the traditionally defined REM sleep onset, and peaked in the minutes right after the transition before decreasing (Figure 3D). Iota activity instead built up more slowly, peaking in the middle of a REM bout, and was sensitive to REM duration (Figure 3E). The different dynamics of beta and iota demonstrate that these should be treated as separate oscillations, and each uniquely qualifies REM sleep.

The specificity of iota to REM sleep is tempered by its presence in wakefulness. Our comparison between iota frequency in REM sleep and in wake reveal that the oscillations are correlated, however not excessively (R=.3), with a bias for wake iota being slightly slower than in REM sleep. Iota in wakefulness on average appears over bilateral parietal areas (Snipes, 2025), and in REM sleep on average over frontal areas (Figure 4G), but some variants bridge the two (Figure 5, (Snipes, 2025)). Something similar occurs for alpha and theta in wake and REM sleep. Alpha is found in both wake and REM sleep in occipital channels (Suppl. Figure 4.5, 4.6), although REM alpha is slightly slower than wake alpha (Cantero and Atienza, 2000; Wong et al., 2026). Theta in wake is usually frontal midline (Snipes et al., 2022), whereas in REM sleep it is both frontal and occipital (Suppl. Figure 4.5). Therefore generally, oscillations in wakefulness and REM sleep appear to be related but are not identical.

The finding that iota peaks are consistent across multiple recordings not only validates the detection method, but also confirms that iota activity is a stable trait, comparable to the other major EEG oscillations (Salinsky et al., 1991; Buckelmüller et al., 2006; Näpflin et al., 2007; De Gennaro et al., 2008; Grandy et al., 2013; Popov et al., 2023). This was true for peak frequency, amplitude, bandwidth, and topography (Figure 2C-E, Figure 5E). Likewise, iota frequencies, amplitudes, and topographies (but not bandwidths) were also more similar in monozygotic twins compared to dizygotic twins or unrelated individuals, indicating these factors are genetically determined, like the other major oscillations (Lykken et al., 1974; van Beijsterveldt et al., 1996; De Gennaro et al., 2002; Smit et al., 2005; Ambrosius et al., 2008; Rusterholz et al., 2018). This means that the consequences of iota heterogeneity along these dimensions, if there are any, will be likewise highly heritable.

Despite iota peak properties being stable across measurements at most a few months apart, all these properties showed significant age-related trends. Iota peaks became less prevalent, slower, and of higher amplitude with age (Figure 4A-C). However, the most substantial change was in bandwidth, which switched from being narrow in children to broad in adults, and split in 20-30 year-olds (Figure 4D). All other EEG measures that changed with age did so gradually (Figure 4, Suppl. Figure 4.1–4.4). Such an abrupt change has, to our knowledge, not been found for any other neurobiological measure, and we do not know what developmental milestone this could reflect. Frontal lobe development is known to continue into young adulthood (Kolk and Rakic, 2022), but all related measures change slowly and relatively little across this decade, including gray and white matter volume (Bethlehem et al., 2022), metabolites (Thomson et al., 2024), connectivity (Sun et al., 2025), and other EEG signals (Favaro et al., 2023; Sun et al., 2023). Likewise, this bimodality in young adults does not appear for phasic REM duration nor overall REM duration (Bertin et al., 2026).

Narrowband and broadband iota may not even be functionally equivalent. Multipolar iota topographies were more often from narrowband peaks and most common in children, while unipolar topographies were more often broadband and in adults (Figure 5), suggesting different source networks. Narrowband iota activity increased across sleep cycles in children, but broadband iota in adults did not (Figure 4H), suggesting broadband iota is less related to REM sleep architecture. The significance of this abrupt change from narrow to broadband iota is completely speculative at this point; it may be as trivial as finding one’s first gray hairs, or as consequential as the legal definition of adulthood.

Aside from age, there was only a small effect of sex on iota peak amplitudes (higher in females), which could be explained by anatomical differences in skull thickness (Lynnerup et al., 2005). There was no relationship to either ADHD or insomnia. According to the framework of insomnia being driven by REM sleep fragmentation (Riemann et al., 2012, 2025; Van Someren, 2021), a link between iota oscillations and insomnia could have been expected, however, a challenge in such group-level comparisons is that the trait-like differences in baseline amplitude may supersede any changes that reflect sleep architecture (Pierson-Bartel and Ujma, 2024). Iota peak amplitudes were correlated by 96% across two nights, potentially masking any differences in REM quality or quantity. It may be more feasible to identify changes in sleep quality through changes in REM iota activity within participants.

While this study used an ideal setup of hundreds of high-density recordings, there are key limitations. First, the combination of the two datasets (Zurich and Amsterdam) divided the population almost exactly into children and adults, with some overlap in the key 20-30 year old range. There were sufficient recordings in this overlapping range confirming that the narrow- to broadband jump was not due to differences in dataset (Suppl. Figure 4.10), but it may have exaggerated or underestimated the difference. Aside from the location and study populations, the datasets differed in experimental protocols, electrode montages, and data quality. All the data is biased towards European participants, and given the strong genetic component, ethnicity effects on iota are likely. Thus, these results may not generalize worldwide. Likewise, the study population did not include children under 7, nor older adults over 70. Wilkinson et al. found broadband 30 Hz oscillations in infants during wake, which may or may not be the precursor of iota (Wilkinson et al., 2024). On the other end of the scale, the finding that all bandwidths increase linearly with age may reflect progressive neurodegeneration, which may accelerate in the last decades of life or in patients with neurodegenerative disorders. Thus, the age related trends observed here may not be neatly extrapolated beyond the 7-70 range. Finally, the two patient groups investigated were “populations of opportunity” rather than the target of investigation; they were under heterogeneous treatment regimes, diagnosed from multiple centers. The non-significant group findings do not exclude that these or any other clinical population could have altered iota oscillations. In general, this paper is only the first step in understanding the demographics of iota oscillations.

In conclusion, iota oscillations are a subtle but genuine signal reflecting REM sleep as well as wakefulness. They behave like other oscillations characteristic of states of vigilance: they show high interindividual variability, high night-to-night consistency, high heritability, and changes with age. Their most unusual characteristic is the jump in bandwidth in young adults, reflecting some yet-to-be determined developmental milestone. Although challenging to measure, iota oscillations offer a new opportunity to investigate REM sleep, brain maturation, and the symphony of neural dynamics.

## 5 METHODS

### 5.1 Data

All data was collected with approval from local ethics committees and in accordance with the declaration of Helsinki. Participants signed informed consent prior to participation. Children < 14 years old had a legal guardian sign consent, and for participants between 14-18 years old, both parent and child signed consent. Data was pooled from multiple studies in two centers, Zurich (CH) and Amsterdam (NL). The individual datasets are specified in Table 1. Separate analyses were performed on data from twins collected in Bern (CH).

**Table 1:**
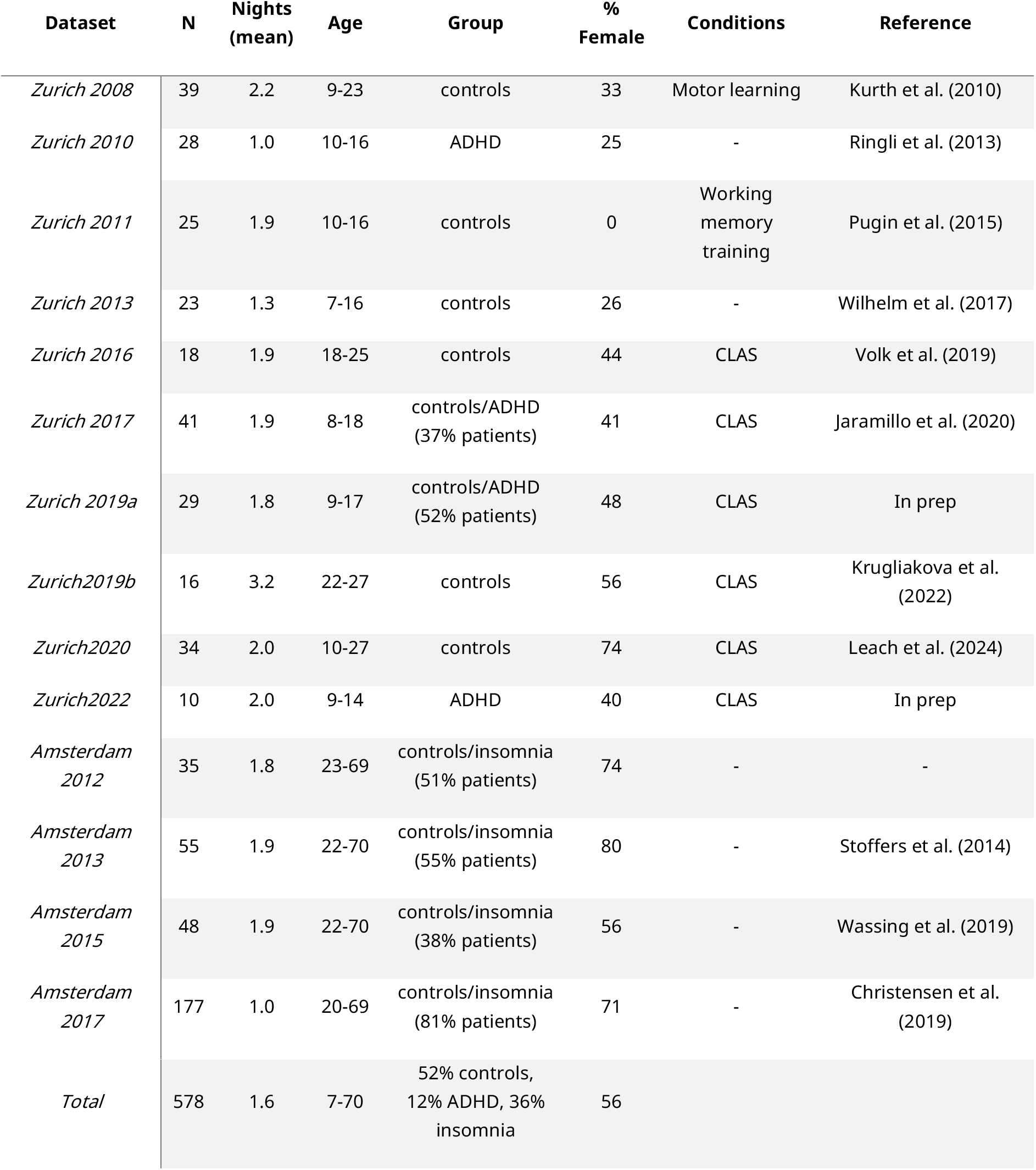
Demographics of the participants included in the analyses of Figure 1 to 5, split by dataset. *Dataset* indicates the start date and city of each study. *N* is the number of participants. *Nights* indicates the average number of recordings each participant had (non-whole numbers reflect repeat measurements for some individuals, dropouts, or recordings lost to data quality). *Conditions* are the experimental conditions applied to each experiment. CLAS: closed loop auditory stimulation. *Reference* papers indicate the main study for a given dataset, or where a majority of participants were previously analyzed. Not all participants were included in the original papers, and not all participants in the original studies were eligible for these analyses (e.g. insufficient REM sleep).

| Dataset | N | Nights (mean) | Age | Group | % Female | Conditions | Reference |
| --- | --- | --- | --- | --- | --- | --- | --- |
| *Zurich 2008* | 39 | 2.2 | 9-23 | controls | 33 | Motor learning | Kurth et al. (2010) |
| *Zurich 2010* | 28 | 1.0 | 10-16 | ADHD | 25 | - | Ringli et al. (2013) |
| *Zurich 2011* | 25 | 1.9 | 10-16 | controls | 0 | Working memory training | Pugin et al. (2015) |
| *Zurich 2013* | 23 | 1.3 | 7-16 | controls | 26 | - | Wilhelm et al. (2017) |
| *Zurich 2016* | 18 | 1.9 | 18-25 | controls | 44 | CLAS | Volk et al. (2019) |
| *Zurich 2017* | 41 | 1.9 | 8-18 | controls/ADHD (37% patients) | 41 | CLAS | Jaramillo et al. (2020) |
| *Zurich 2019a* | 29 | 1.8 | 9-17 | controls/ADHD (52% patients) | 48 | CLAS | In prep |
| *Zurich2019b* | 16 | 3.2 | 22-27 | controls | 56 | CLAS | Krugliakova et al. (2022) |
| *Zurich2020* | 34 | 2.0 | 10-27 | controls | 74 | CLAS | Leach et al. (2024) |
| *Zurich2022* | 10 | 2.0 | 9-14 | ADHD | 40 | CLAS | In prep |
| *Amsterdam 2012* | 35 | 1.8 | 23-69 | controls/insomnia (51% patients) | 74 | - | - |
| *Amsterdam 2013* | 55 | 1.9 | 22-70 | controls/insomnia (55% patients) | 80 | - | Stoffers et al. (2014) |
| *Amsterdam 2015* | 48 | 1.9 | 22-70 | controls/insomnia (38% patients) | 56 | - | Wassing et al. (2019) |
| *Amsterdam 2017* | 177 | 1.0 | 20-69 | controls/insomnia (81% patients) | 71 | - | Christensen et al. (2019) |

**Table 1: Demographics of the participants** included in the analyses of Figure 1 to 5, split by dataset.
|  |  |  |  |  |  |
| --- | --- | --- | --- | --- | --- |
| *Total* | 578 | 1.6 | 7-70 | 52% controls,<br>12% ADHD, 36%<br>insomnia | 56 |

#### 5.1.1 Zurich Children’s Hospital Dataset

Data from 306 participants were recorded from the Children’s Hospital of Zurich, between 7 and 26 years old, collected between 2008 and 2022. After excluding participants for insufficient clean REM sleep, the final dataset included 263 individuals. Of these, 68 were diagnosed with ADHD (9-17 years old, 19 female) by the Department of Child and Adolescent Psychiatry and Psychotherapy at the University of Zurich, the outpatient clinic of the Child Development Center, and at private children’s clinics in Oerlikon, Zurich. ADHD patients were not excluded based on medication status and therefore were a mixture of medicated, previously medicated, and unmedicated (see (Furrer et al., 2019)). All participants were screened for good sleep quality, general health, no (other) medication, and no (other) comorbidities.

The majority of recordings were done investigating the effects of closed loop auditory stimulation (CLAS) during NREM sleep (Volk et al., 2019; Jaramillo et al., 2020; Krugliakova et al., 2022; Leach et al., 2024). Other recordings either had no experimental manipulation (Ringli et al., 2013; Wilhelm et al., 2017), involved a motor learning task prior to sleep (Kurth et al., 2010; Wilhelm et al., 2014), or a 3 week working memory training (Pugin et al., 2015). The experimental conditions were not investigated in this analysis.

Sleep time was determined by participants’ individual preferred sleep and wake-up time, which they had to maintain the week prior to each measurement. All data was recorded using 128-channel Electrical Geodesics Inc (EGI) Geodesic Sensor gel-based nets and EGI amplifiers. Recordings were done with Cz reference, either at 500 or 1000 Hz sampling rate, impedances set to below 50 kOhm at sleep onset.

All data was low-pass filtered at 50 Hz using EEGLAB’s *pop_eegfiltnew()* function (a zero-phase Hamming-window FIR high-pass filter, with automatically detected filter orders), then downsampled to 200 Hz to reduce file size. External channels 49, 56, 107, 113, and cheek channels 126 & 127 were excluded from all analyses.

Sleep was scored by trained experts using standard AASM scoring guidelines (Berry et al., 2012), on 20 s epochs.

#### 5.1.2 Netherlands Institute for Neuroscience Dataset

Data from 349 participants was made available from the Netherlands Institute for Neuroscience (18-70 years old), recruited between 2012-2022, pooling multiple studies (Table 1) of either the Free University of Amsterdam Medical Center or University of Amsterdam (Stoffers et al., 2014; Wei et al., 2017; Christensen et al., 2019; Wassing et al., 2019; Cox et al., 2024). Of these, 315 had sufficient clean REM sleep epochs, for a total of 442 recordings. 209 participants were diagnosed with insomnia disorder. Healthy controls had to have no reported sleep difficulties and an Insomnia Severity Index (ISI) score < 8. All participants were excluded for sleep apnea, restless leg syndrome, narcolepsy or other somatic, neurological or psychiatric disorders, use of sleep medication within the prior 2 months, and irregular circadian rhythms.

256 channel high-density EEG was recorded with EGI Geodesic gel-based sensor nets (1000 Hz sampling rate, Cz reference). Electrode impedances were kept below 100 kΩ. Edge and cheek channels were removed (82, 91, 92, 102, 111, 120, 133, 145, 165, 174, 187, 199, 208, 209, 216, 217, 228, 229, 233, 235:243, 245, 246, 247, 251, 255, 256, 258:265), and remaining channels were interpolated to correspond to the same locations as the 128 net configuration (using EEGLAB’s *pop_interp()*). Data was then further low-pass filtered and downsampled to 200 Hz.

Sleep was scored by trained experts using AASM guidelines on 30 s epochs.

#### 5.1.3 Data subsets

Depending on the analysis, different subsets of the datasets were used. Sample sizes (N) or degrees of freedom (df) are provided for each statistic accordingly.

An initial 1131 recordings were made available for this study. Of these, 937 had at least 10 minutes of clean REM sleep (epochs with no more than 20 artefactual channels) and were included in the analysis. Recordings could be excluded due to damaged/incomplete files, too many artefacts (see section 5.2), or too little REM sleep.

The final dataset included 578 participants 7 to 70 years old, 56% female. For quantifying interindividual differences, e.g. related to age, one night per participant was selected. When multiple nights were available, the night with the most REM sleep in minutes was chosen.

For analyses of prevalence (e.g. mixed effects models, proportion of recordings) the entire dataset of multiple nights per participant was used (N = 937). When comparing night-to-night differences, only participants with at least two nights were used (N = 313). When comparing iota properties, only individuals with an iota peak were included (N = 349 for REM sleep), or only the subset with peaks > 0.30 (N=174). The latter was done for example when comparing transitions into and out of REM sleep (Figure 3D), when the data was z-scored therefore giving equal weight to individuals with low or high amplitude activity.

#### 5.1.4 Bern Twins Dataset

A separate EEG dataset from 62 twins (MZ = 18, DZ =13) aged 9-15 years old (MZ: 12.8 ±1.3 y.o.; DZ: 12.9 ± 1.4 y.o.) was analyzed. This dataset was previously published (Inderkum and Tarokh, 2018; Markovic et al., 2018, 2020, 2022; Rusterholz et al., 2018). Participants were healthy, good sleepers.

Data was recorded on a 64 electrode (58 EEG channel) gel-based EGI Geodesic system, recorded in the participants’ homes, with 1000 Hz sampling rate, Cz reference. Prior to recordings, participants spent at least 5 days on a stabilization schedule, ensuring 9.5-10 h of sleep. Caffeine intake was prohibited for 3 days before and during the study.

Sleep was scored using AASM guidelines on 30 s epochs.

Data was already processed before being reused for this analysis (Markovic et al., 2018). EEG data was high-pass filtered with a 10 s median sliding window, low-pass filtered at 100 Hz, then downsampled to 250 Hz, and average referenced. Power density spectra were calculated for each 30 s epoch using Welch’s method (5 s Hanning windows, no overlap).

### 5.2 Preprocessing: artefact rejection and spectral power calculation

Analyses were performed with MATLAB v2025b and EEGLAB 2025.0. Analysis code for this manuscript is available at: https://github.com/snipeso/iota-demographics. The preprocessing library is at: https://github.com/snipeso/sleep-prep. Spectral parametrization (MATLAB implementation of specparam): https://github.com/snipeso/eeg-oscillations.

Data was analyzed in “segments” of 20 or 30 s to align with sleep scoring, separately for each channel (simultaneous segments from all channels is referred to as the “epoch”). Artefacts were detected and spectral power calculated for each segment, and power during artefactual segments were excluded from analyses.

Artefact rejection was done automatically. Artefacts were initially detected in the time domain (see section 5.2.1). The Cz channel was restored and then EEG data was re-referenced to the average, excluding artefactual segments. Spectral power was calculated for each average-referenced segment (section 5.2.2). Specparam was applied to each spectrum, the output of which was used for a second artefact rejection in the spectral domain (section 5.2.3). The twin data only used the latter artefact rejection, as it was provided already as spectral power.

Segments were additionally removed if they were ≤ 1 minute between artefacts (such gaps could contain sub-threshold artefacts). Epochs were removed if more than 20 segments were artefactual, or if any segment did not have at least 2 clean neighbors (this did not apply to edge segments). Channels with less than 30 min of clean data were likewise marked entirely as artefactual. Remaining artefact segments were set to NaN and were not included in analyses, but did not limit the minimum clean data requirements when averaging segments.

#### 5.2.1 1st artefact rejection: time-domain

Time-domain artefacts were detected for each sample point in four categories:

1. Large-amplitude artefacts, where timepoints were flagged when the signal exceeded ±1000 µV or changed by more than 100 µV between consecutive samples (padded by 1 s).
2. Flat lines, in which the moving standard deviation of a window of 10 s fell below 0.1 μV.
3. Disconnected channels, when 5 or more electrodes picked up identical environmental noise (30 s segments correlated by ≥ .999), or when a segment didn’t correlate with more than 5 channels by more than .3.
4. Muscle artefacts were detected by filtering the EEG between 20–45 Hz (*pop_eegfiltnew*), calculating the power envelope (squared Hilbert amplitude), and smoothing it with a 0.2 s moving mean. Bursts exceeding 100× the median power across all channels and timepoints were flagged as artefacts. Artefact edges were extended to include adjacent timepoints where power remained above 20× the same median.

For the above artefact detection, a separate copy of the data was high-pass filtered at 0.5 Hz with *pop_eegfiltnew()*, and notch filtered (Kaiser-window FIR filter; 48–52 Hz stopband, 2 Hz transition bands, automatically determined order using *keiserord* and zero-phase forward–backward filtering using *filtfilt*) to remove 50 Hz line noise. Artefacts 1-3 were first detected with Cz reference, then large artefacts were set to 0, just for the purpose of muscle artefact detection, since these artefacts can propagate to all channels during averaging. The data was average-referenced, and then muscle artefacts were detected.

Artefact labels were resampled to the scoring epoch resolution (20 or 30 s), such that any segment with even a single artefactual timepoint was marked as an artefact.

#### 5.2.2 Spectral power and spectral parametrization (specparam)

Power spectral density (PSD) was computed on 20 or 30 s segments using Welch’s method (MATLAB function *pwelch*; window = 4 s Hanning, overlap = 50%). The power spectrum was smoothed prior to spectral parameterization using a 3-Hz moving median filter followed by a 2-Hz mean LOWESS filter (explained in Suppl. Figure 4.11A).

The aperiodic (1/f) component was separated from the periodic component using the specparam algorithm (formerly FOOOF; (Donoghue et al., 2020)), implemented in MATLAB (Snipes et al., 2026). The frequency range was between 2-45 Hz, peak width = 0.5-20 Hz, minimum peak height 0.1, unrestricted number of peaks, fixed mode. Otherwise specparam defaults were used. The outputs include: periodic power (reflecting oscillatory activity), aperiodic exponents (slope of the background signal), aperiodic offsets (overall spectral amplitude), model fit measures of mean absolute errors (MAE) and R^2.

This study quantifies spectral activity as periodic power, either averaged within a given band or as the peak’s amplitude. This is in log10 space, unitless after subtracting the aperiodic component. Throughout the paper, the unit of measurement is indicated as “pp” (periodic power) for disambiguation.

This calculation per segment was done for the preprocessing (section 5.2.3), but the same steps were applied to epoch-averaged data for the main analyses to extract periodic power.

#### 5.2.3 2nd artefact rejection: spectral-domain

Segments were also excluded based on specparam outputs: MAE > 0.15, R² < 0.95, aperiodic exponent outside 0.5 to 4.5, or aperiodic offset outside −1 to 5 (Suppl. Figure 4.12A-D). Segments were further excluded if their exponent deviated from the remaining epoch’s median by > 0.6. The latter usually affected single channels with unusually high high-frequency noise (Suppl. Figure 4.12A), but also captured the loss of signal quality occurring gradually across the night in some recordings, possibly from the electrode gel drying up. Lastly, remaining segments were removed if the relationship between exponent and offset deviated from the otherwise linear trend present in the data by |z| > 10 first within, then across channels (Suppl. Figure 4.12E-F).

#### 5.2.4 (The absence of) ICA artefact rejection

For these analyses, unlike in Snipes (2025), we did not perform independent component analysis (ICA) to remove physiological artefacts (heartbeat, muscle, and eye movements). Given the size of the dataset (>900 recordings), half of which were of 256 electrodes (ICA must be performed prior to interpolation), this would have been computationally intensive and time consuming.

We used one of the datasets (Zurich 2020) to develop the current preprocessing pipeline and tested the utility of ICA when detecting iota activity. We found that the primary benefit was in removing heartbeat components which affect the iota (but especially beta) range in edge channels (Suppl. Figure 4.11C,D). We chose to exclude edge channels when identifying spectral peaks and found this solution sufficient for our current analyses (although it hindered the manual classification of weaker iota peak topographies; Suppl. Figure 5.1). However, future investigations of high frequency activity in REM sleep, especially involving beta activity, would benefit from also performing ICA.

### 5.3 Analysis

#### 5.3.1 Channel averaged spectral peaks

To create an overview of periodic activity during each sleep stage, without prior assumptions of spectral bands (Figure 1), we applied the analysis in Snipes (2025). Untransformed PSD from clean segments were averaged across channels except the edge channels ([1, 8, 14, 17, 21, 25, 32, 38, 43, 44, 48, 57, 63, 64, 68, 69, 73, 74, 81, 82, 88, 89, 94, 95, 99, 100, 114, 120, 121, 125, 128]), and across epochs for each sleep stage. Then the specparam algorithm (section 5.2.2) was applied to extract periodic power for each sleep stage. Stages with less than 10 min of artefact-free epochs were excluded.

Unlike in Snipes (2025), which used the specparam output for periodic peak properties, here spectral peaks were identified in a second step from periodic power using the MATLAB function *findpeaks* (MinPeakProminence = .01 pp, MinPeakDistance = 1 Hz), providing the amplitude and center frequency of each peak. Peak bandwidth was then calculated as the distance to the first frequency (either before or after the center frequency) that reached 50% of the peak amplitude, multiplied by 2. This is an approximation of the full-width to half-maximum (FWHM), taking into account that iota peaks could sometimes be found half-merged with beta peaks (e.g. Figure 3A), and therefore using only the most conservative estimate of bandwidth. Because the previous iota analysis (Snipes, 2025) relied on the peaks outputted by specparam directly, “bandwidth” was then defined as 2 times the standard deviation of fitted gaussians to each peak, which are slightly narrower than the FWHM described here.

Peaks were discarded if another larger peak was within 1/4 of its bandwidth. Harmonics were further removed whenever they occurred within ±1 Hz of the second or third harmonic of a stronger lower-frequency peak.

We established the edges of the iota bands, separately for wake and REM sleep, by calculating the mean and standard deviation of iota peak frequencies of our sample population, then modelling the population-level normal distribution, and considered a band which would encompass 95% of this fitted distribution. First, peak frequencies detected between 2-45 Hz were pooled across participants within each stage. Their distribution was smoothed using kernel density estimation (MATLAB’s *ksdensity*), and local maxima were identified using *findpeaks*. These maxima represent the frequencies at which detected oscillatory peaks were concentrated. For iota, we selected the maximum closest to 30 Hz and defined preliminary boundaries using the nearest local minimum on either side or, if no minimum was present, the point where density fell below 5% of the maximum. A normal distribution was then fitted to peak frequencies within these boundaries using *fitdist*. Final band boundaries were defined as the fitted mean ± 2 standard deviations. As a reference check, applying the same procedure to wake peaks near 10 Hz yielded the expected alpha range of 8-12 Hz.

#### 5.3.2 Maximum spectral peaks

To increase sensitivity for iota peaks that were too localized to appear in channel averages, we developed the following peak detection approach to select the maximum peak in a given sleep stage for a given band across all channels. First, untransformed PSD from all segments of a given sleep stage and channel were averaged (minimum 10 min of epochs per stage), then specparam was applied with the same processing and parameters in section 5.2.2 to the average spectra of each channel.

For the maximum peak to be considered a true peak, it had to have at least 4 other channels that had a peak within 0.5 Hz of the same frequency, and at least 50% of the same amplitude. Candidate peaks within a band were tested in descending amplitude order; when a peak failed these criteria, the next candidate peak was tested until one passed or none remained. This could exclude spurious peaks such as those due to both environmental noise as in Suppl. Figure 2.2B, and unusual muscle activity as in Suppl. Figure 2.2F. While the latter was rare in REM sleep (due to muscle atonia), in NREM sleep edge channels could sometimes show spectral peaks in 1-2 channels, with little consistency even in neighboring channels, indicative of artefacts.

The following bands were used: theta 4-8 Hz, alpha 8-12 Hz, sigma 12-16 Hz, beta 16-25 Hz, iota 25-40 Hz. Delta (0.5-4 Hz) was not included because a) the aperiodic signal was measured starting from 2 Hz and is generally unreliable at the edges; b) most traditional “slow wave activity” in the delta range is captured by the aperiodic signal (Bódizs et al., 2021). 40-50 Hz gamma was not included because few peaks were detected in this range (Figure 1). Since the data was collected in Europe, data around 50 Hz was not considered due to line noise.

Given that harmonics were not a substantial contributor to REM sleep iota (Suppl. Figure 1.2), potential harmonics were not excluded from analyses like was done for Figure 1 to avoid excluding iota peaks that spuriously were a multiple of a theta or alpha peak. Thus, iota in NREM in Figure 3C and supplementary figures also include sigma harmonics; regardless, these harmonic peaks were small enough not to obscure the dominance of iota activity during REM sleep, especially by using custom bands based on REM sleep iota.

#### 5.3.3 Custom band power

When investigating spectral power in individual REM bouts or sleep cycles, or timecourses such as transitions into REM sleep, custom bands were identified based on the previously described peaks in the average spectra, as peak frequency ± bandwidth/2. Then, periodic power (already calculated for each segment from the artefact detection procedure) was averaged in each band, providing a timecourse for each custom band. Epochs with artefacts were excluded. Band power was then averaged for the top 10 channels expressing the highest REM peak topography (averaged across all epochs). If no REM sigma peak was identified, the N2 sigma peak was used.

#### 5.3.4 REM band transitions

Transitions into and out of REM sleep were quantified by fitting sigmoids to the previously described band timecourses. First, the timecourses were smoothed with a 1 min moving mean. Missing intervals of ≤ 1 min were filled with linear interpolation. The resulting timecourses were then linearly up-sampled to a common 5 second resolution.

Transition windows were selected taking 5 min prior and 5 min after transitions into and out of REM sleep. For transitions into REM sleep, only REM bouts of 5 min or longer were considered, with no REM sleep in the 5 min prior (likewise for the 5 min after REM in transitions out of REM sleep). Any band’s transition with > 1/3 epochs marked as artefacts was excluded. Within each night, all timepoints of transitions into REM sleep were z-scored for each band (and again for transitions out of REM sleep). Then, all transitions into REM sleep were averaged within a participant, pooling multiple recordings.

While the previous approach allowed an exploratory visualization of the transition (Figure 3D), to calculate onset times for each band, the transition was further simplified. The data was padded before and after the transition by 10 min. The timepoint with the lowest value in the 5 min before the transition was taken as the start of the transition, and all values before that was assigned this minimum value (including the padded data). Likewise, the highest value in the 5 min after the transition was assigned to all values from that timepoint onward. Then, an increasing sigmoid was fitted to the data (MATLAB central file exchange *sigm_fit*). The opposite was done for transitions out of REM sleep. The choice of padding ensured comparable fit stability for slow and faster transitions. The extension of the lowest and highest point took into account the initial surge of beta at transition onset observed in Figure 3D. Restricting the analysis to increasing sigmoids at REM onset and decreasing at REM offset excluded participants who displayed the opposite dynamics (e.g. beta activity increasing more during wake after REM offset, which occurred when beta power was stronger in subsequent wake, 10-20% of recordings). Excluding such cases did not bias estimating transition timing. Sigmoid fits in which the transition point was outside ± 5 min from the onset were likewise excluded.

The sigmoid fit provides four parameters: starting and ending values, the midpoint of the transition in time, and the speed of the transition. Analyses focused on the midpoint of the transition.

#### 5.3.5 REM band trajectory, within bouts

To quantify how each band behaves within REM sleep, the same smooth custom-band timecourses as above were used, again including only REM bouts that were 5 min or longer, in Figure 3D, middle. Each bout of varying length was then averaged into 10 bins, providing normalized time. This data was then z-scored within a recording, then averaged across all REM bouts of each participant.

#### 5.3.6 REM measures per sleep cycle

REM duration was quantified within sleep cycles for Figure 3E and mixed effects models. Sleep cycles were calculated by first assigning the initial wake period to N2, and then all subsequent wake and N1 bouts to whichever sleep state was previously scored. N2 and N3 were merged, thus simplifying sleep staging to just NREM and REM sleep. Each cycle began at the onset of a NREM period lasting >10 minutes and ended immediately before the next qualifying NREM period. REM duration was the total amount of REM in minutes during each cycle.

#### 5.3.7 Topography type classification

Scoring topography type was done manually by author SS, randomizing and blinding recordings (Suppl. Figure 5.1). Each topography could be scored as either: frontal bilateral, frontal unilateral, frontal midline, parietal bilateral, parietal unilateral, quadripolar, pentapolar, tripolar up point, tripolar down point, or undetermined. Ten categories were chosen out of convenience (keys 0-9), but different subtypes could be merged or separated, fairly arbitrarily. A topography was scored as unilateral when a second pole could not be visually identified, but in practice bilateral topographies were often partially lateralized, forming a continuum with completely lateralized types. Quadripolar types encompassed also topographies with only 3 peaks but these were placed in the typical lateralized frontal and parietal locations. Undetermined topographies were either fairly uniform without any clear peaks, or artefactual channels on the edges were the strongest signal. Occasionally poor data quality resulted in splotchy topographies. The presence of one or multiple peaks in the spectra was simultaneously scored manually. Eventually it will be possible to automate classification, but this was not deemed a good use of time presently. Manual classification further allowed verification that artefacts were not driving the iota peak selection.

### 5.4 Statistics

All statistics were done with α = .05. Correction for multiple comparisons was done with False Discovery Rate (FDR) correction (Benjamini and Hochberg, 1995).

When comparing two distributions, within subjects paired t-tests were conducted (t_paired), or between subjects unpaired t-tests (t_unpaired). Effect sizes are provided as Cohen’s d, evaluated such that values around 0.2 are considered small, 0.5 medium, and 0.8 or above large (Cohen, 1988). When comparing the relationship between two continuous variables, Pearson’s correlations were conducted (R).

To evaluate the relationship between two variables while also controlling for various factors and interactions with other variables, linear mixed effects models were used, each model provided in proximity to the reported results. Random intercepts were used for participants, and individual recordings when appropriate. The full model output is provided as a supplementary file. For evaluating the measure of iota peak prevalence, binomial generalized linear mixed models were used.

To evaluate measurement consistency, intraclass correlation coefficients (ICC[A,1]) were used, and evaluated according to the criteria of (Koo and Li, 2016), whereby < .5 is poor, .5-.75 is moderate, .75-.9 is good, and .9-1 is excellent.

Night-to-night iota topography consistency was assessed by conducting Pearson’s correlations of custom periodic power across all channels to two nights within participants. These were compared to the correlations between participants with a one-sided permutation test (10,000 permutations).

Twin data was analyzed descriptively with ICC and Pearson’s R. Within-pair similarity of REM iota peak frequency, amplitude, and bandwidth was quantified separately in monozygotic (MZ) and dizygotic (DZ) twins using a one-way random-effects, single-measure ICC(1,1), and was reported with 95% confidence intervals. Frequency and bandwidth analyses were restricted to pairs in which both twins had a detected peak. For amplitude analyses, amplitude was set to 0 when REM data were available but no iota peak was detected.

Heritability of peak variables was assessed with structural equation modelling (SEM), previously applied to this dataset (Rusterholz et al., 2018; Markovic et al., 2022), but here implemented in MATLAB. Variance was partitioned into additive genetic (A), shared environmental (C), and unique environmental and measurement-error (E) components. ACE, AE, and CE models were fitted by maximum likelihood, and the model with the lowest AIC was selected.

The heritability of iota topographies was first evaluated descriptively by comparing the correlations across channels within MZ and DZ twins using a two-sided Welch’s t-test.

For this test, R-values were Fisher-z transformed. Heritability was further quantified using Falconer’s estimate, h2 = 2(rMZ-rDZ), with shared environmental contributions estimated as c2 = 2rDZ-rMZ. Topographies were calculated over each participant’s custom iota-frequency band, and only participants with a detected peak were included.

## ACKNOWLEDGEMENTS

This project was funded by The Swiss National Science Foundation Postdoc.Mobility Grant (P500-3_235448) and the European Research Council (ERC) Starting Grant SleepingAwake (101116748).

The original datasets were collected with funding provided by the Swiss National Science Foundation (PP00A-114923, PP00P3-135438, CRSII3_160803, 320030_153387, 320030_179443, PCEFP1_181279, 2003B_184943, 10.003.398), the University Research

Priority Program of the University of Zurich, the Clinical Research Priority Program “Sleep and Health” of the University of Zurich, the HMZ Flagship Project “SleepLoop” of University Medicine Zurich, the Borbély-Hess Foundation, Gottfried und Julia Bangerter-Rhyner-Stiftung, Jacobs Foundation, the Waterloo Foundation (2462-4978), the Bial Foundation grant 252/12, the Netherlands Organization of Scientific Research (NWO) grant VICI-453.07.001, and the ERC (ERC-ADG-2014-671084 INSOMNIA, ERC-ADG-2020-957641 INSOMNIA AID, and ERC-ADG-2021101055383 OVERNIGHT).

We’d like to thank Melanie Furrer, Mirjam Studler, Maya Ringli, Renato Merki, Michelle Suppiger, Léa Geiger, Martina Liechti, Stefano Maurizio, Dr. Renate Drechsler, Stephanie Cares, Margot Schaerer, Jessica Bühler, Natalie Birnbaum, Demy Marcella Gutjahr, Cécile Abati, Tamara Weil, Paulina Schmiedeberg, Nicole Meier, Vanessa Kasties, Veronika Voskresenska, Gila Norup, Soraya Souissi who contributed to data collection.

Thank you also to Henry Hebron, Alejandro Osorio-Forero and Maria Dimitriades for feedback and discussions on EEG data analysis, and Simone Accascina for feedback, computing resources, equipment, and technical support.

Claude Sonnet and Opus, as well as ChatGPT Codex (up to v6.1 Sol) were used to assist with some code writing and minor manuscript editing. All AI written code and text was reviewed and edited by a human expert.

## Contributions

### Current investigation

SS: Conceptualization, Methodology, Software, Formal analysis, Visualization, Writing – original draft, reviewing & editing. RC, EK, SK, SL, AM, VJ, CV, FP, IW, LT, EvS, RH: Writing – review & editing. TA: Supervision, Funding acquisition, Writing – review & editing.

### Original investigations from which data were derived

RC: Data curation. EK, SK, SL, AM, VJ, CV, FP, and IW: Conceptualization, Investigation, Methodology, Project administration, Funding acquisition. LT, EvS, and RH: Conceptualization, Methodology, Resources, Supervision, Funding acquisition.

## Conflicts of interest

R.C. is employed by and holds a financial interest in Deep Sleep Technologies B.V. The company was not involved in the design, execution, interpretation or funding of the present study.

## Supplementary Figures

**Suppl. Figure 1.1:**
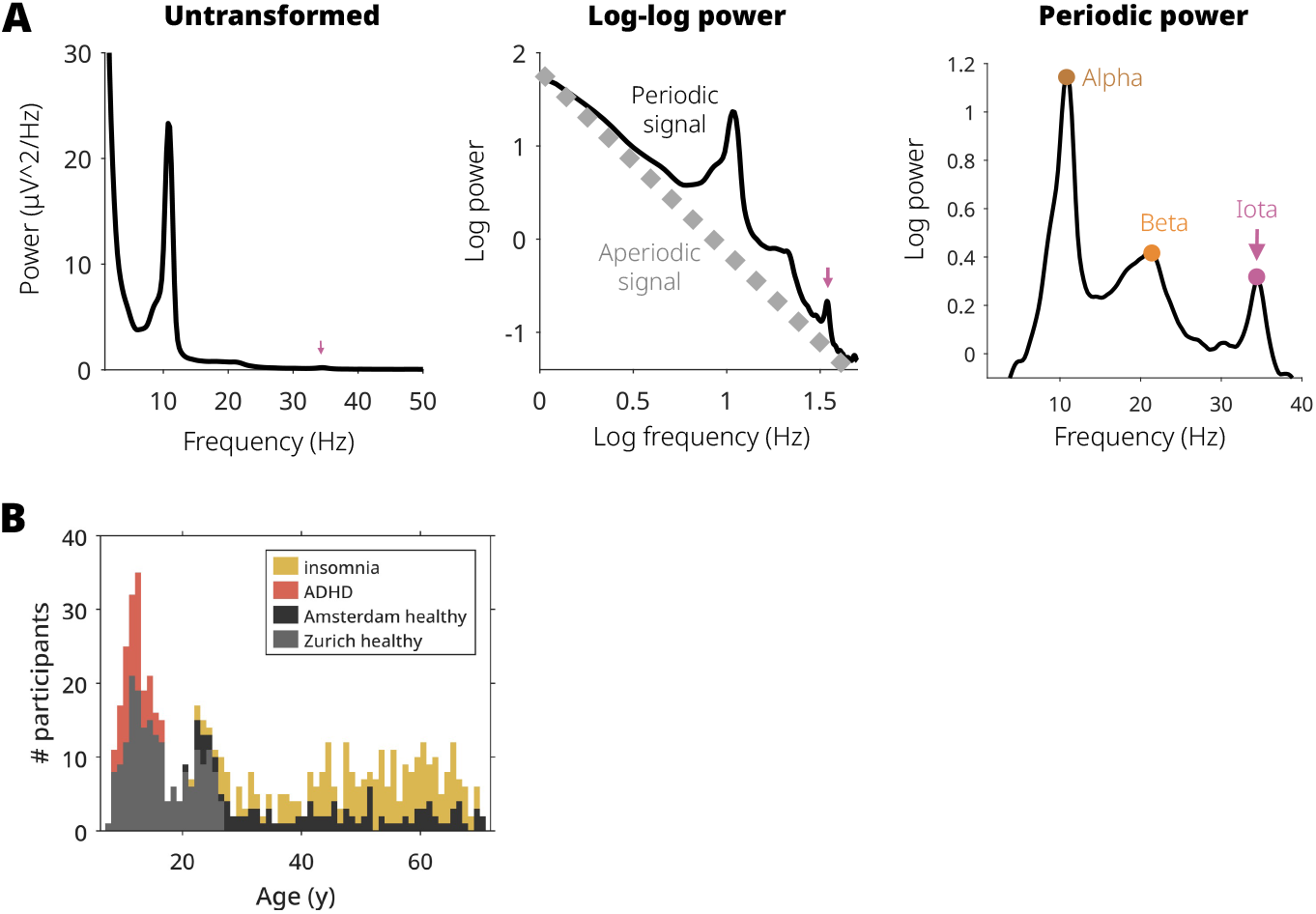
Methods. A) Specparam transformation of spectral power to detect periodic peaks. Example data from wake. Note that the iota peak is almost imperceptible in the untransformed data. B) Stacked histogram of participants, color coded by dataset and patient status.

**Suppl. Figure 1.2:**
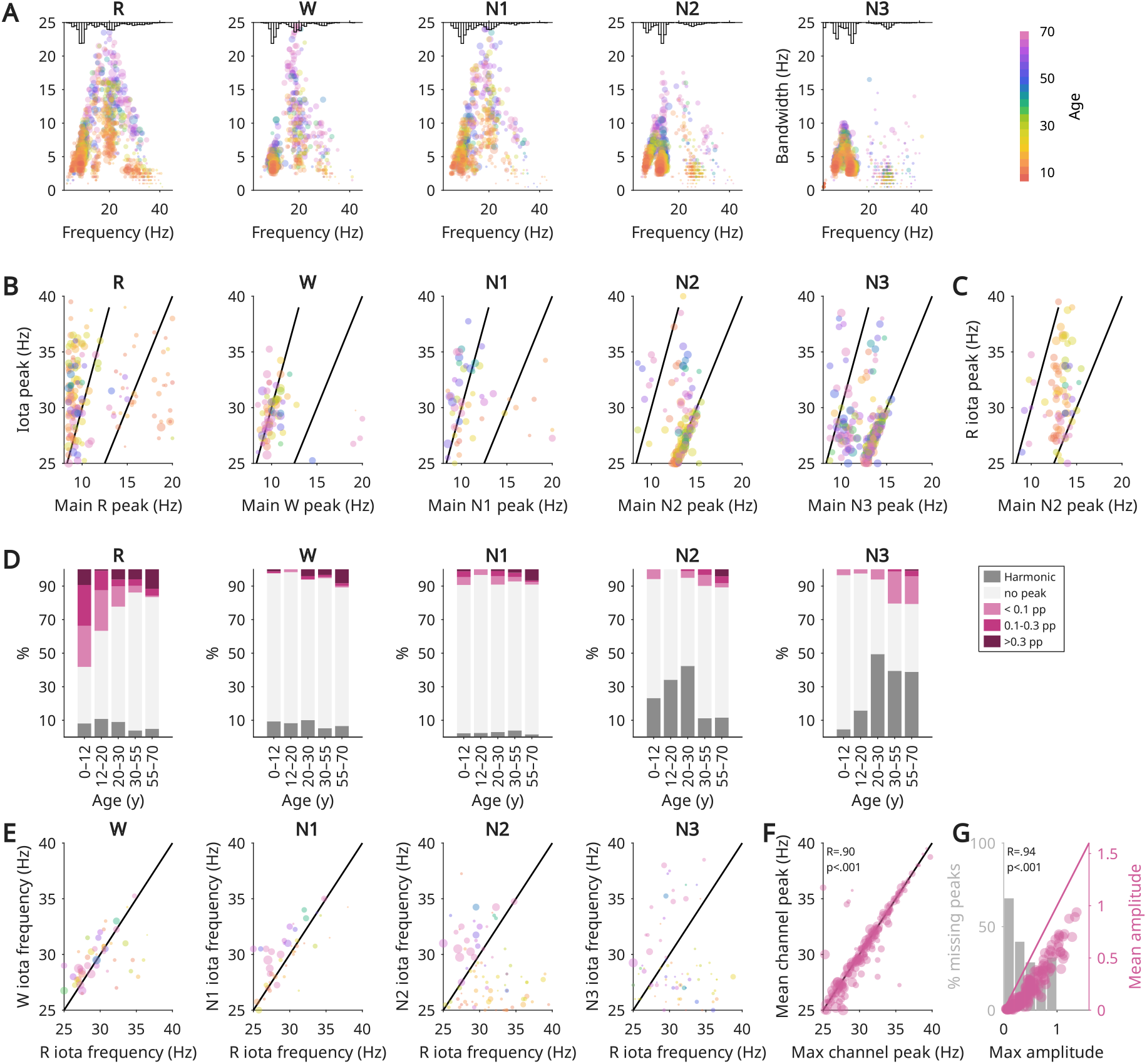
Iota harmonics across sleep stages. Like in Figure 1, here peaks are identified in average EEG channels. A) Same as Main Figure 1, except all peaks, including potential harmonics, are included. Spectral power was averaged within each sleep stage across channels, and all peaks were detected. The x axis is the peak frequency, the y axis is the peak’s bandwidth, the size of each dot reflects the peaks’ amplitude and the color is the age of the participant. Note the small cluster in the high-beta low-iota range in N2 and N3, absent in Main Figure 1. B) Relationship between the highest amplitude oscillation between 8-20 Hz, and that between 25-40 Hz within each sleep stage. C) Relationship between N2 sigma peak and REM iota peak. D) Proportion of iota peaks detected which were multiples of a lower frequency peak (dark gray) or not (pink). The majority of iota peaks detected in NREM were harmonics of sigma. In wake, the distribution is evenly split, and so it is less evident whether iota peaks in wake are harmonics of alpha. There appear to be age effects of harmonics, with N2 iota more likely to be a harmonic in children and young adults, and N3 iota more likely to be a harmonic in adults. E) Correlation between REM sleep iota peaks and every other stage’s iota peaks. Wake and N1 iota are significantly related to REM sleep iota, more so than peaks during NREM sleep. F) Correlation between REM sleep iota peak frequencies detected using channel averages vs. taking the maximum peak of all channels. G) Correlation between amplitudes detected using channel averages or maximum peak of all channels. Predictably, channel-average peaks were systematically smaller, and missed ∼40% of peaks identified from the maximum channel.

**Suppl. Figure 2.1:**
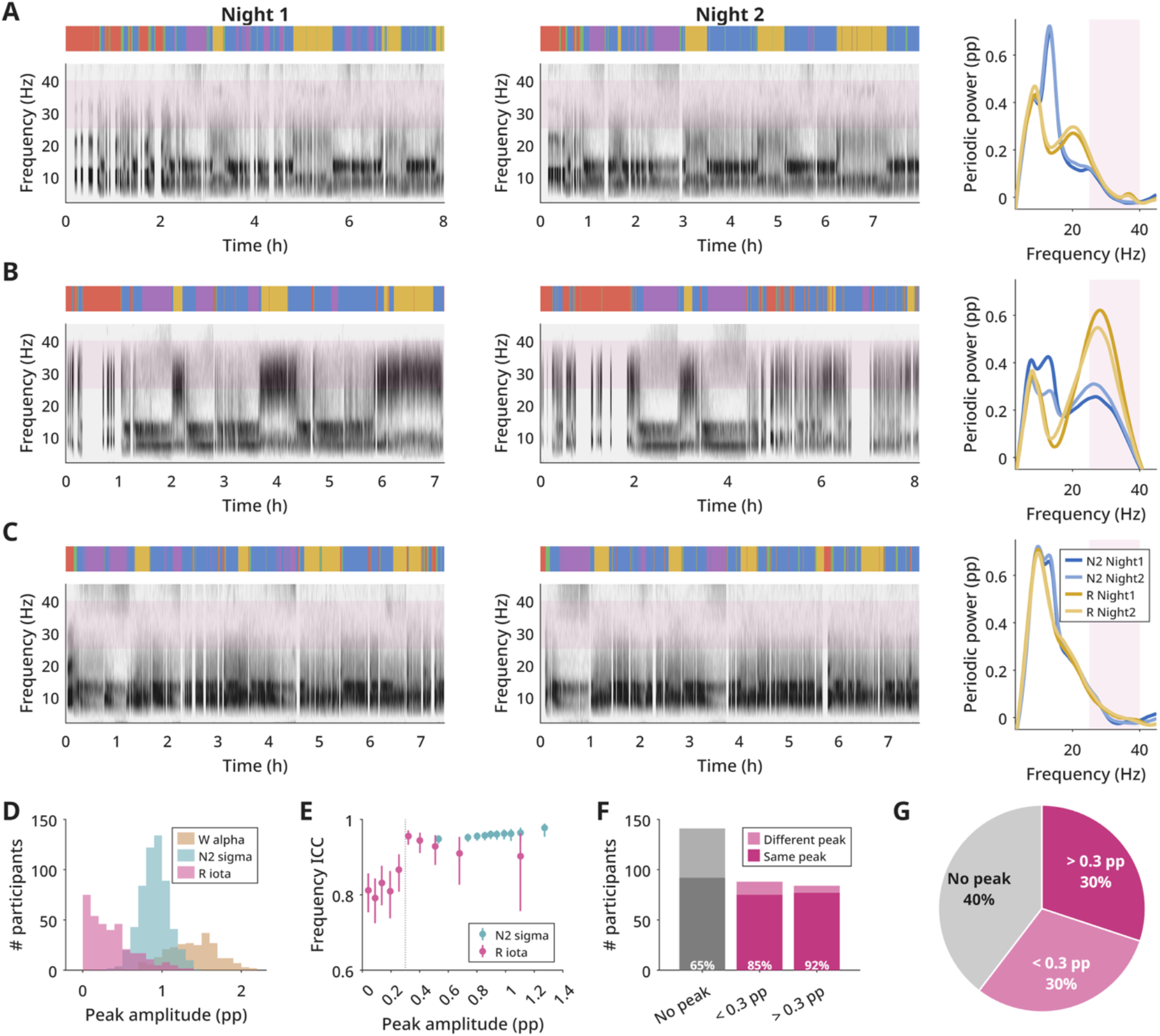
A-C) Examples of the heterogeneity of iota activity. Each row is one participant. Left and middle columns are two nights for the same individual. Each night contains a multi-colored bar plotting AASM manual sleep staging (red=wake, green=N1, blue=N2, purple=N3, yellow=REM). Below it is the spectrogram of periodic power for each epoch, channel-averaged, with darker colors indicating higher amplitudes. A pink band highlights the iota range. The right-most column is the channel- and epoch-averaged spectra for N2 and REM sleep, from each night. A is a low-amplitude iota peak, B is a broadband iota peak, C is a recording with no peak. D) Distribution of peak amplitudes for wake alpha, N2 sigma and REM iota, across all participants. E) Intraclass correlation coefficients (ICC) of iota frequency across two sessions for increasing quantiles of peak amplitudes. Each dot is the ICC for a given quantile of participants and all quantiles larger. The vertical line is the 95% confidence intervals. F) number of participants who had peaks within 1.5 Hz across two sessions (same peak) or not, split by average peak amplitude. ‘No peak’ indicates the number of participants for which no peak was detected in either session (dark gray) or in only one session (light gray). G) Proportion of all recordings within each peak amplitude range.

**Suppl. Figure 2.2:**
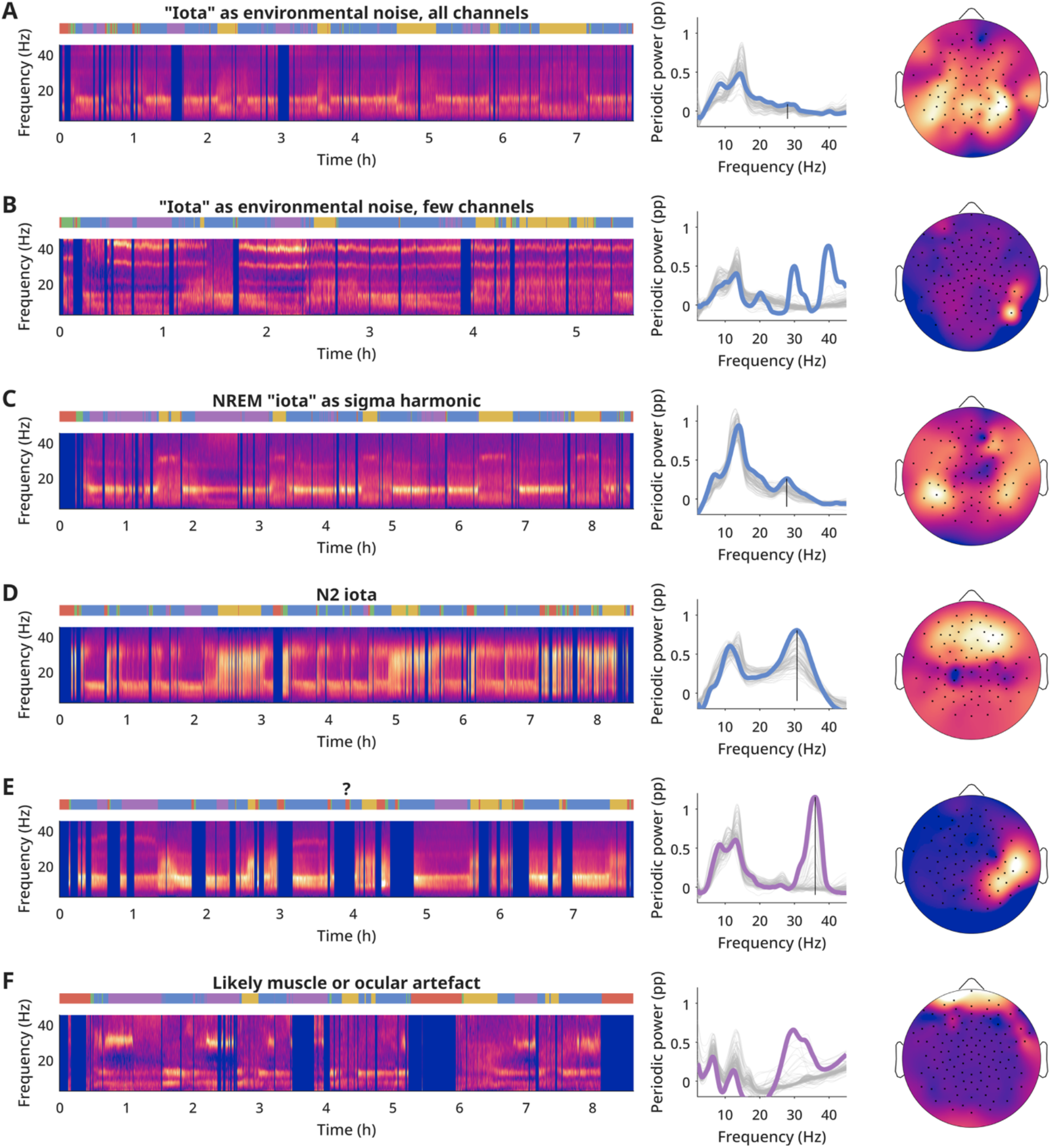
Iota during NREM sleep, representative examples. Left column: time-frequency periodic power and AASM scoring as in Main Figure 2 (red = wake, green = n1, blue = n2, purple = n3, yellow = REM). Middle column: epoch-averaged NREM power. Gray lines are from each channel, the thick line is the channel with the highest amplitude iota peak. Blue indicates the spectra are from N2, purple from N3. Vertical line indicates the iota peak. If the line is absent, no iota peak was identified because the candidate peak did not fulfill all the prerequisites (e.g., not enough channels with similar amplitude and frequency). Right column: epoch-averaged topography for the iota peak. A-B) external noise, equally affecting all sleep stages. C) Iota as a harmonic of sigma. Note: the topography is slightly more lateralized than traditional sigma, as not all sigma sources produce a harmonic. D) Iota during N2, distinct from iota during REM sleep, with a physiological frontal topography. E) “Iota” intermittently appearing in N3 and to a lesser extent N2. Unknown source. F) “Iota” appearing intermittently in both N2 and N3, with poor peak consistency across channels (middle panel), and located at edge channels (right panel), indicating likely a muscle or ocular artefact.

**Suppl. Figure 4.1:**
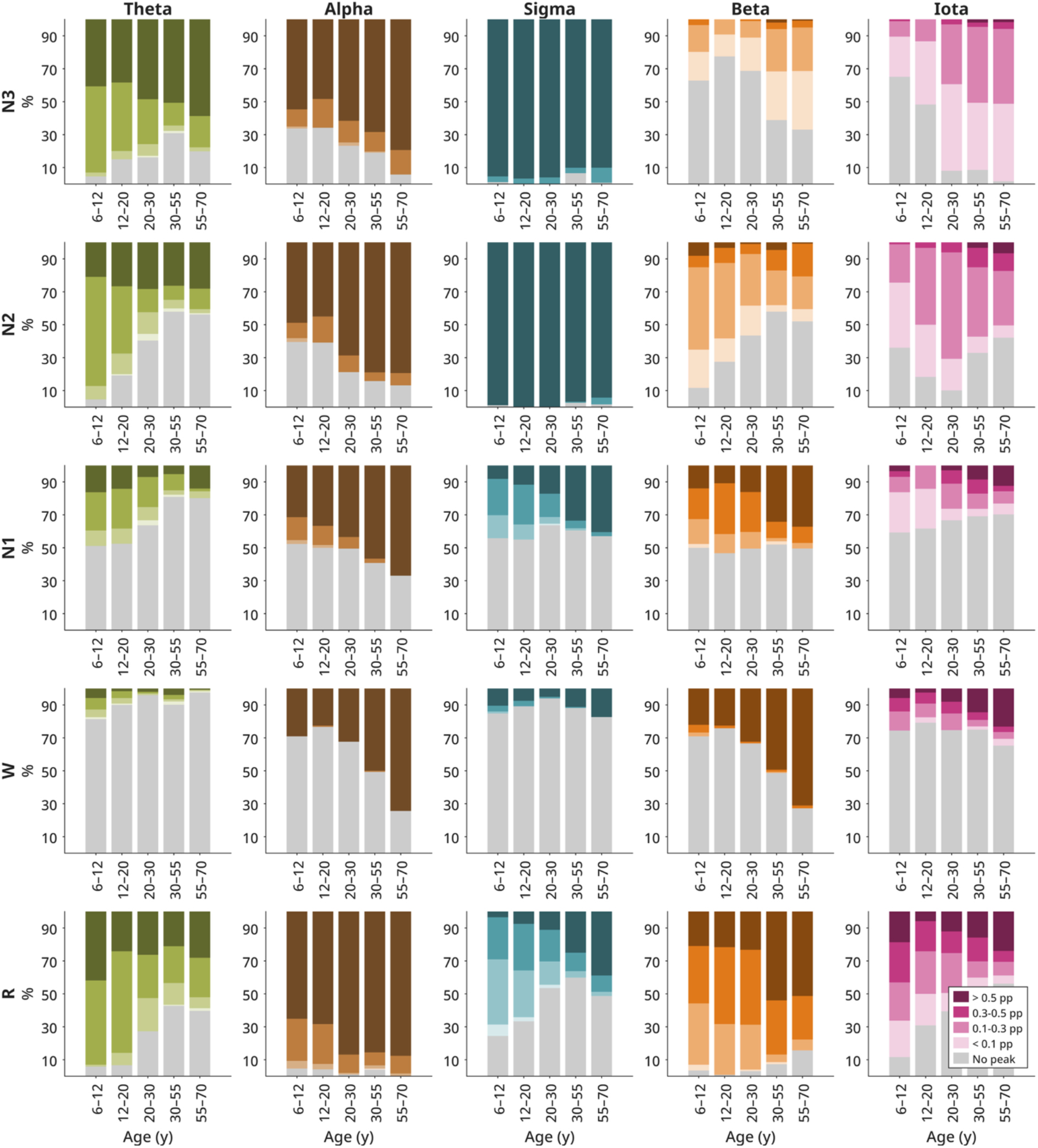
Distribution of detected peaks by amplitude, across ages, same as Main Figure 4A. Note, the iota peaks detected in NREM mostly reflect harmonics (Figure 1.2).

**Suppl. Figure 4.2:**
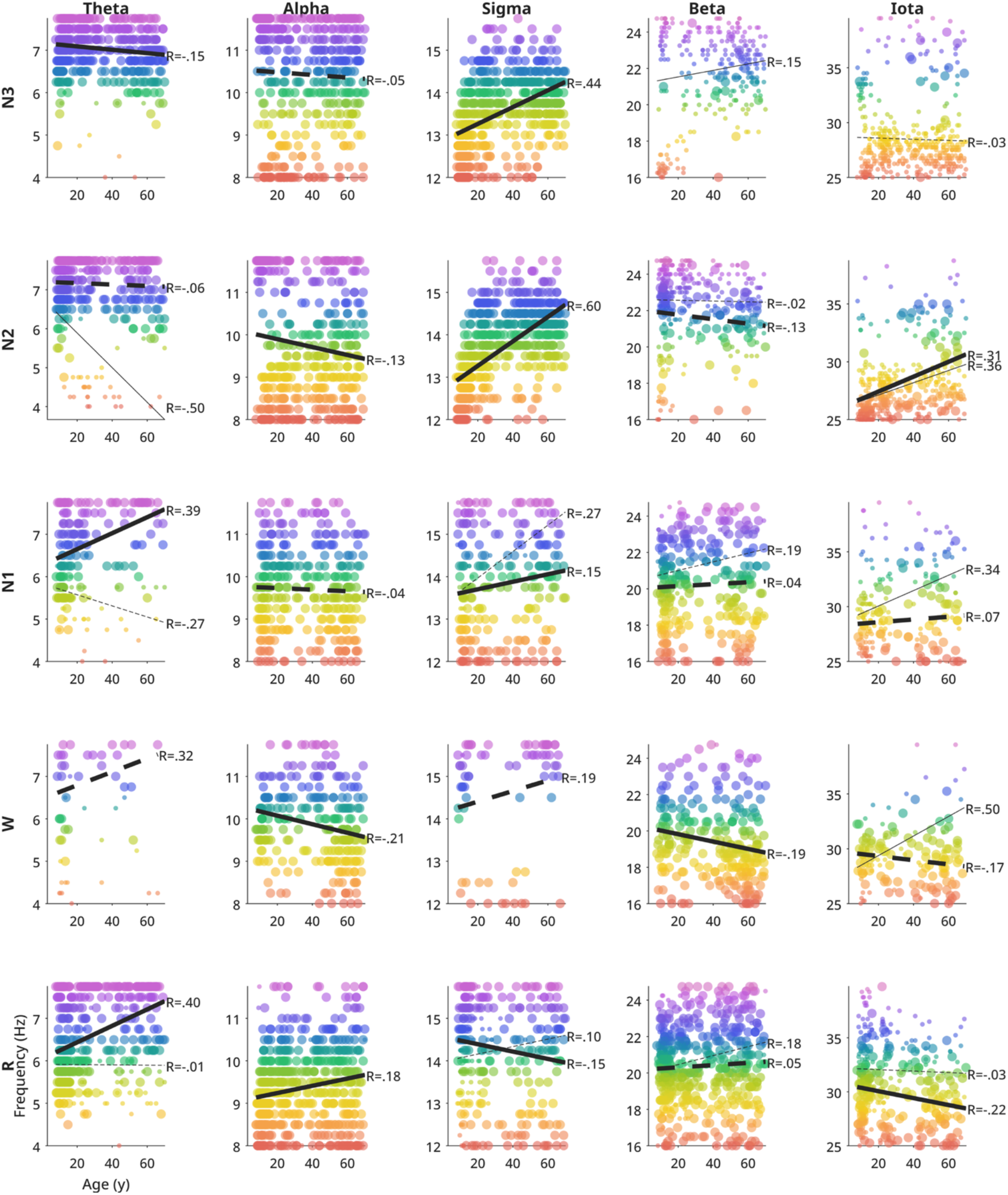
Peak frequencies for all bands and all stages, same as Figure 4B. Color indicates frequency, size indicates amplitude (large dots for peaks > 0.3 pp). One night per participant is included. Thick line indicates linear trend for large amplitude peaks, thin for low amplitude peaks. R values are provided for each line. If the line is dashed, the model was not significant. Note, the standard bands are not always appropriate for sleep, especially across all ages.

**Suppl. Figure 4.3:**
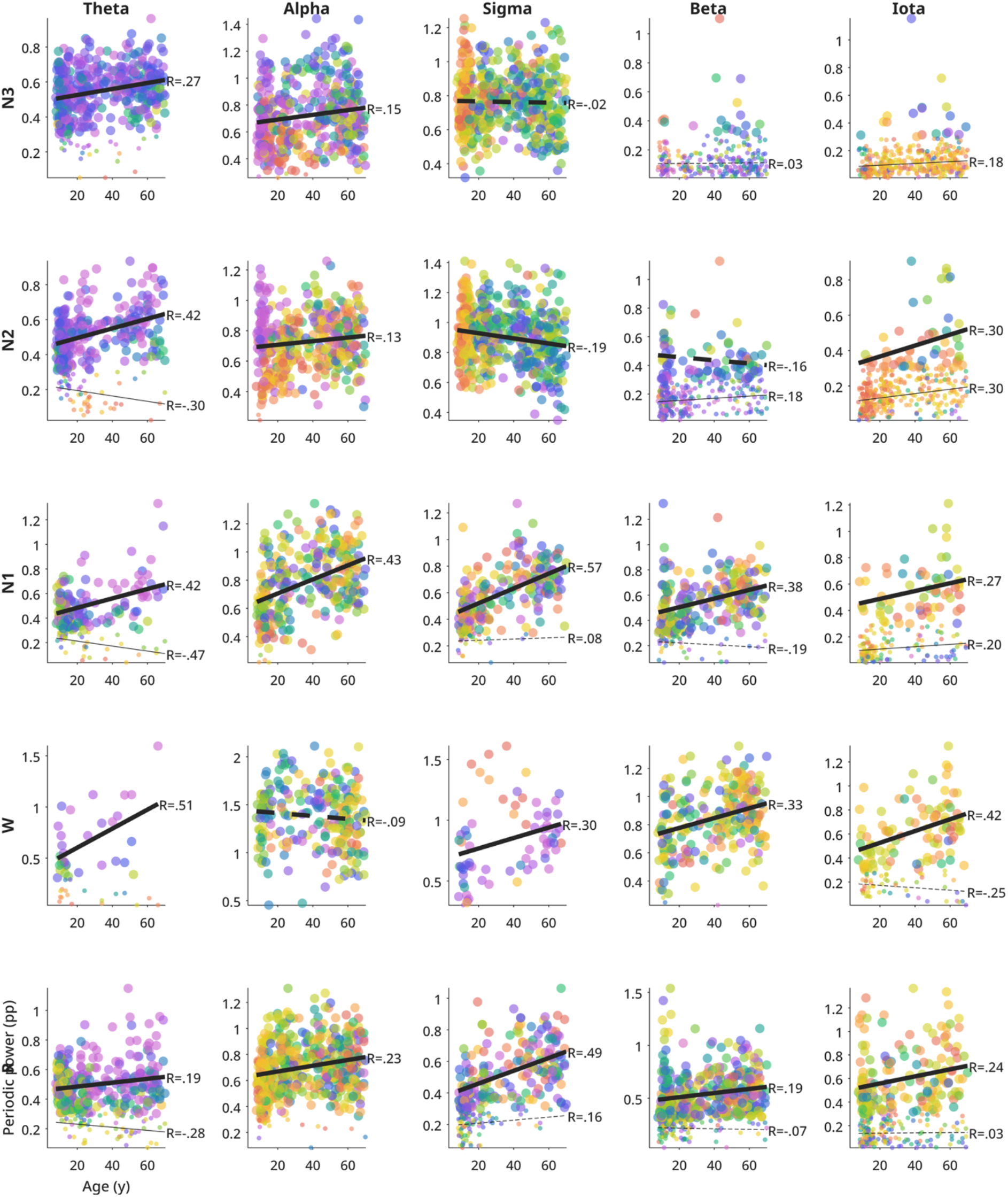
Peak amplitude for all bands and all stages. Same as Figure 4.2.

**Suppl. Figure 4.4:**
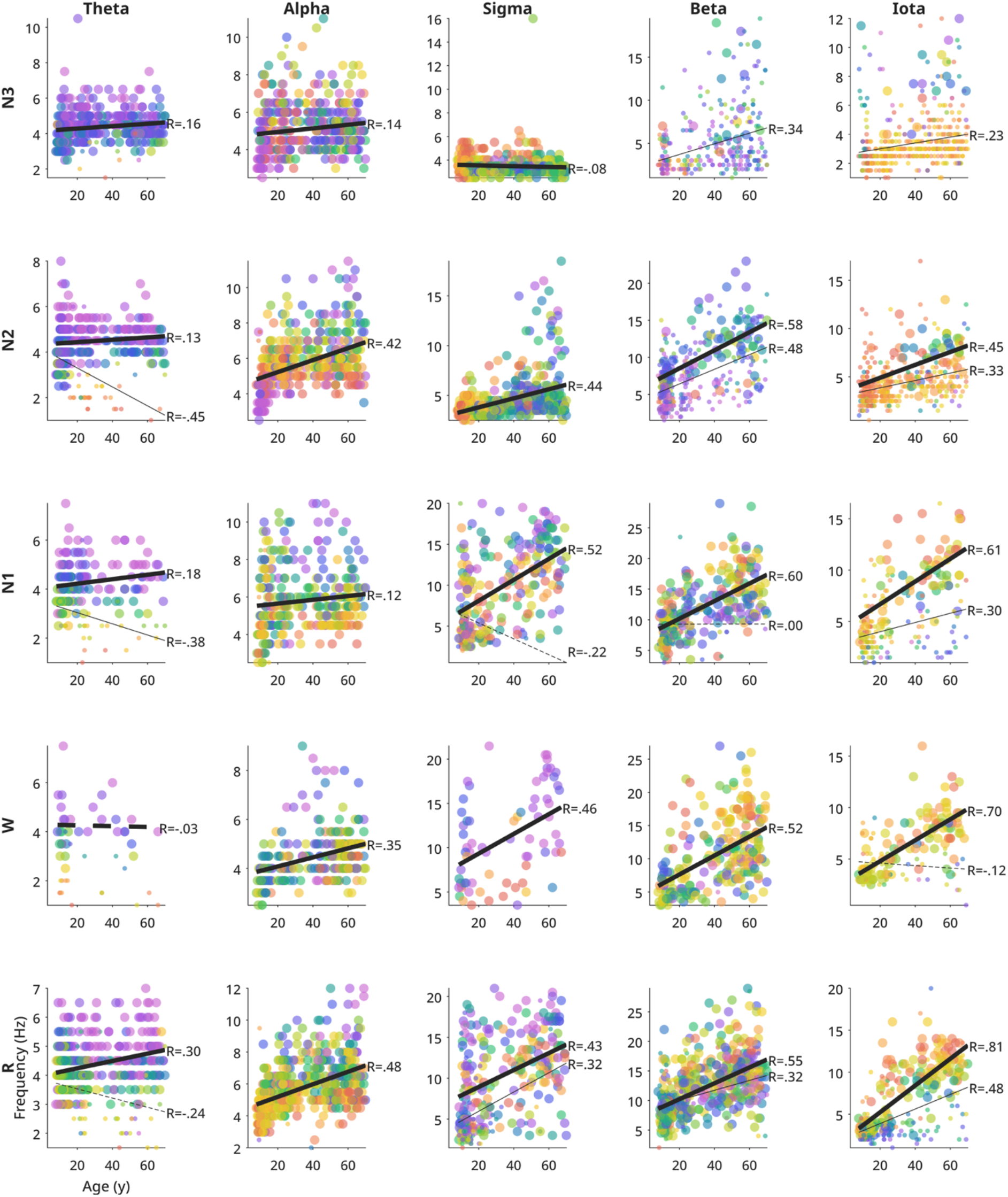
Peak bandwidth for all bands and stages. Same as Figure 4.2.

**Suppl. Figure 4.5:**
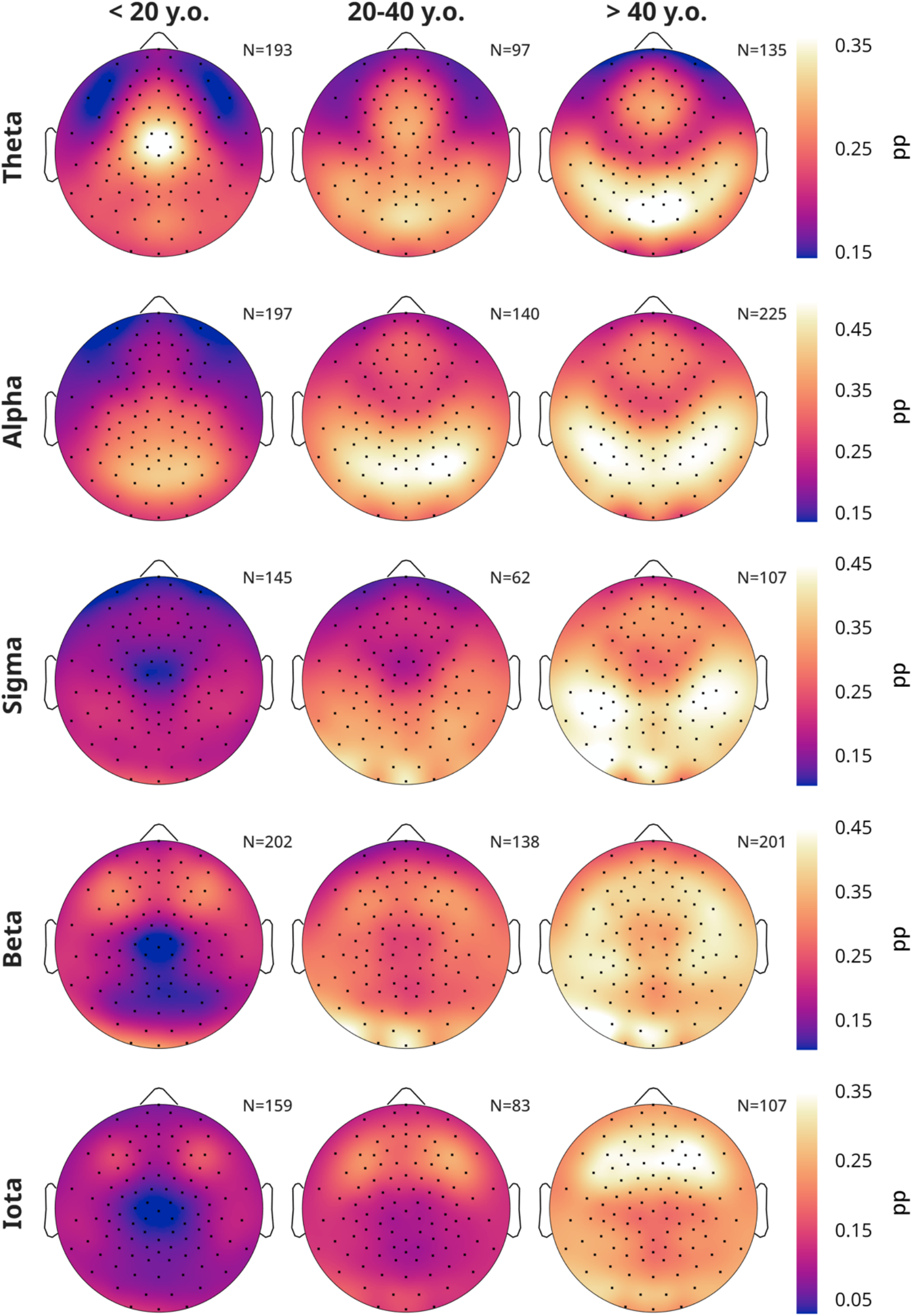
Average R topography per band, split by age. Number of participants included is written in the top right of each figure. Note, the bottom left sources of sigma and beta match that of heartbeat artefacts (Suppl. Figure 4.11C).

**Suppl. Figure 4.6:**
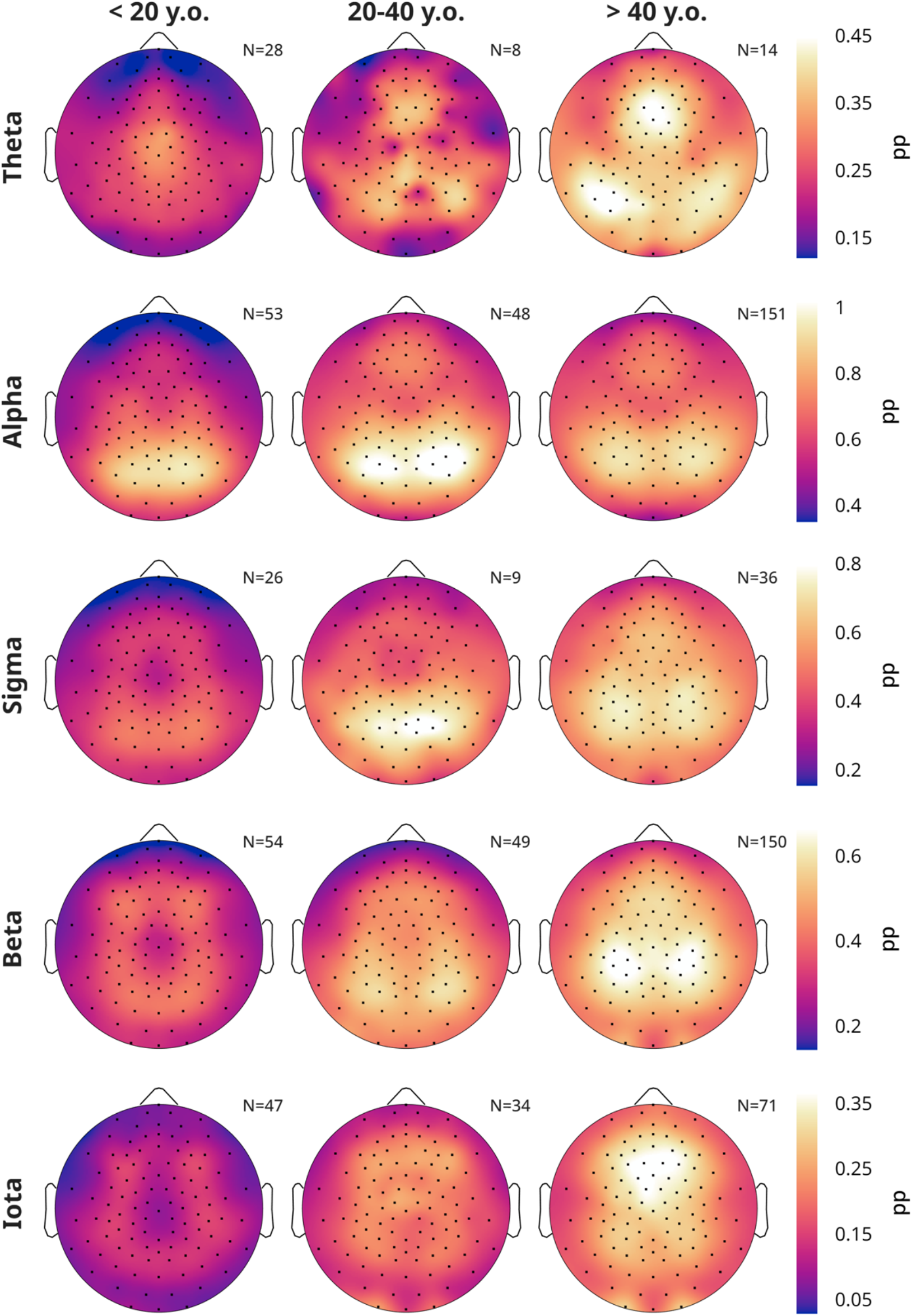
Average W topography per band, split by age.

**Suppl. Figure 4.7:**
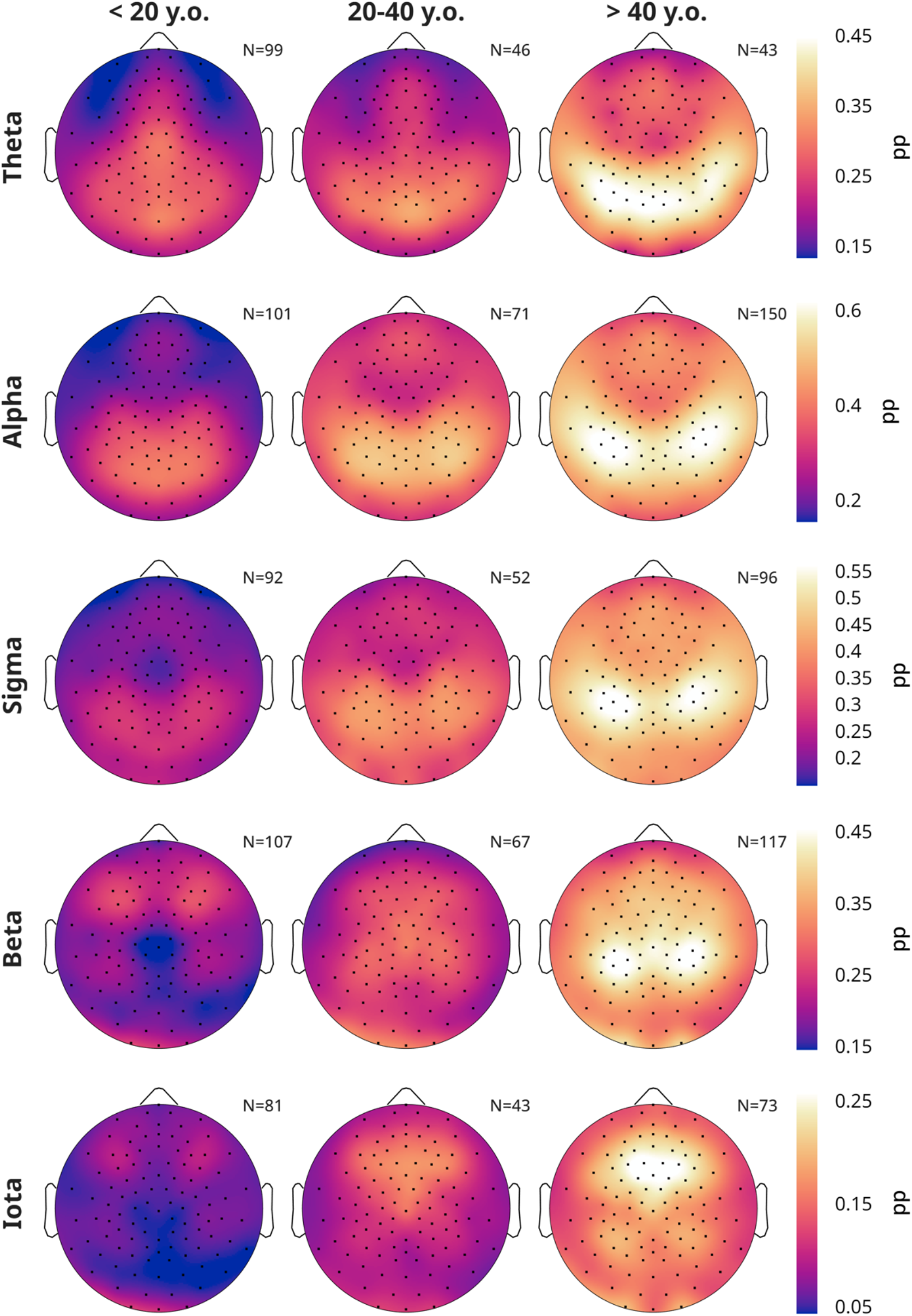
Average N1 topography per band, split by age.

**Suppl. Figure 4.8:**
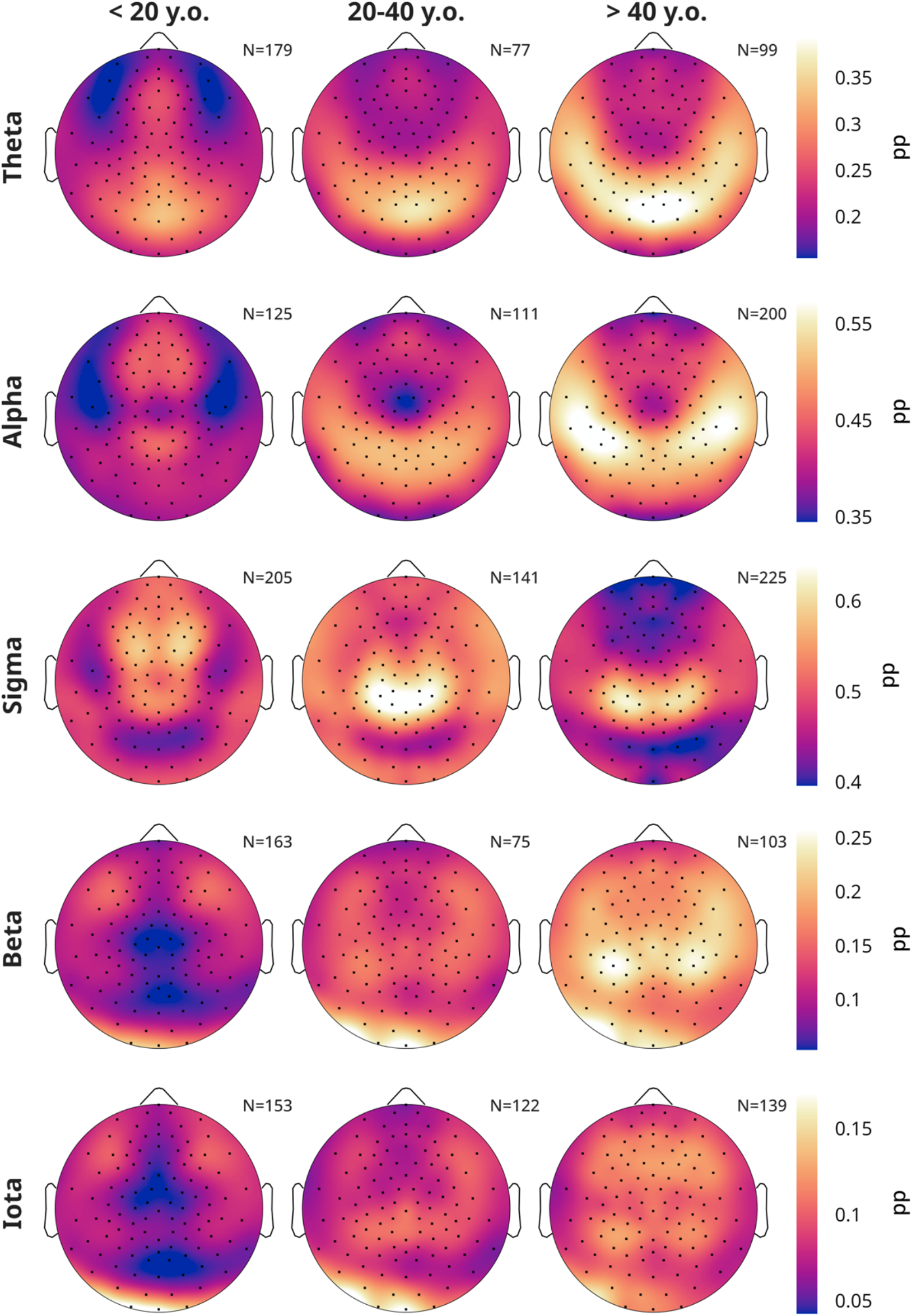
Average N2 topography per band, split by age.

**Suppl. Figure 4.9:**
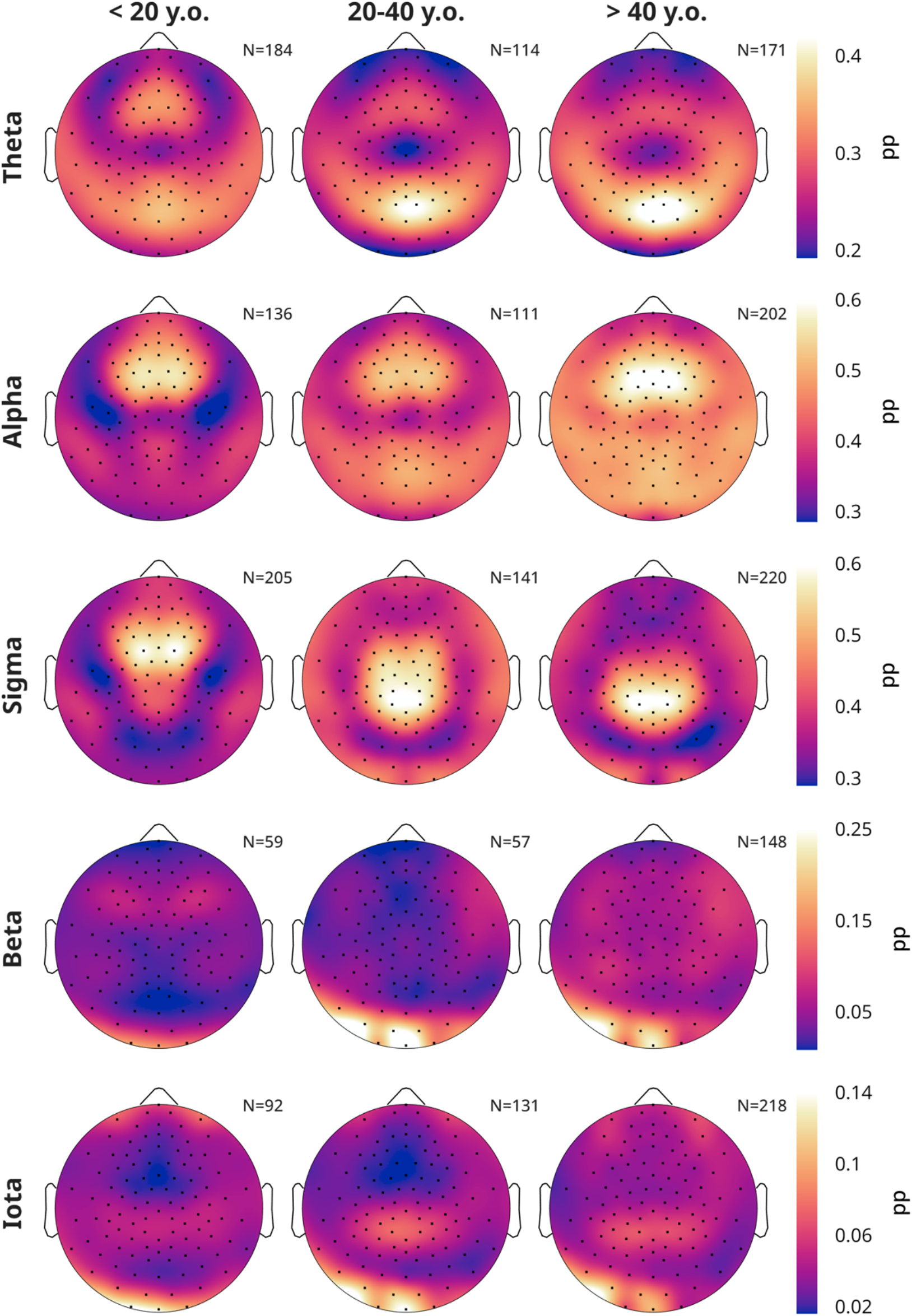
Average N3 topography per band, split by age. Note, ICA artefact rejection was not performed; the bottom edge left cluster in beta and iota reflects heartbeat artefacts.

**Suppl. Figure 4.10:**
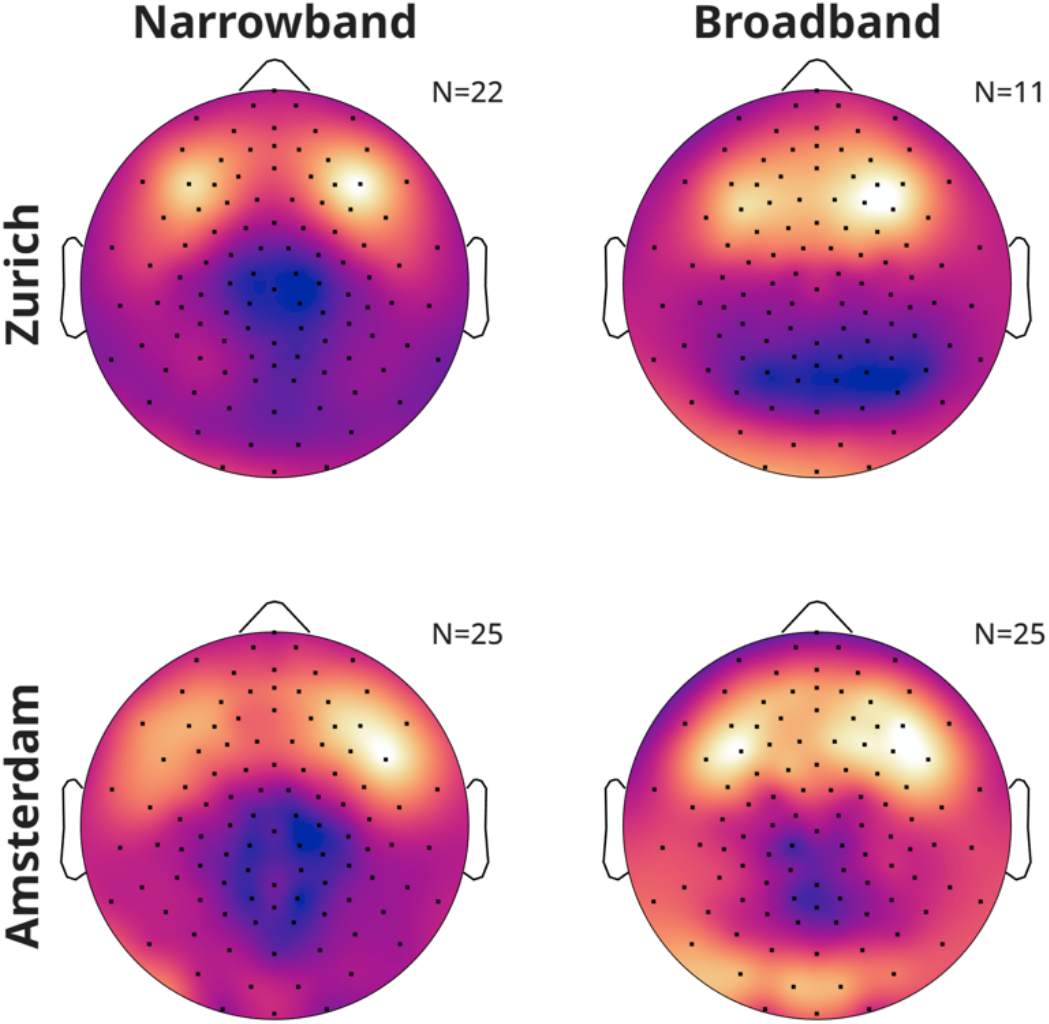
Average REM sleep iota topographies separating recordings from each research center, split by peak bandwidth (more or less than 6 Hz) for participants 20-40 years old. The Amsterdam data has more ovaloid peaks, likely due to the different EEG net montage and subsequent interpolation.

**Suppl. Figure 4.11:**
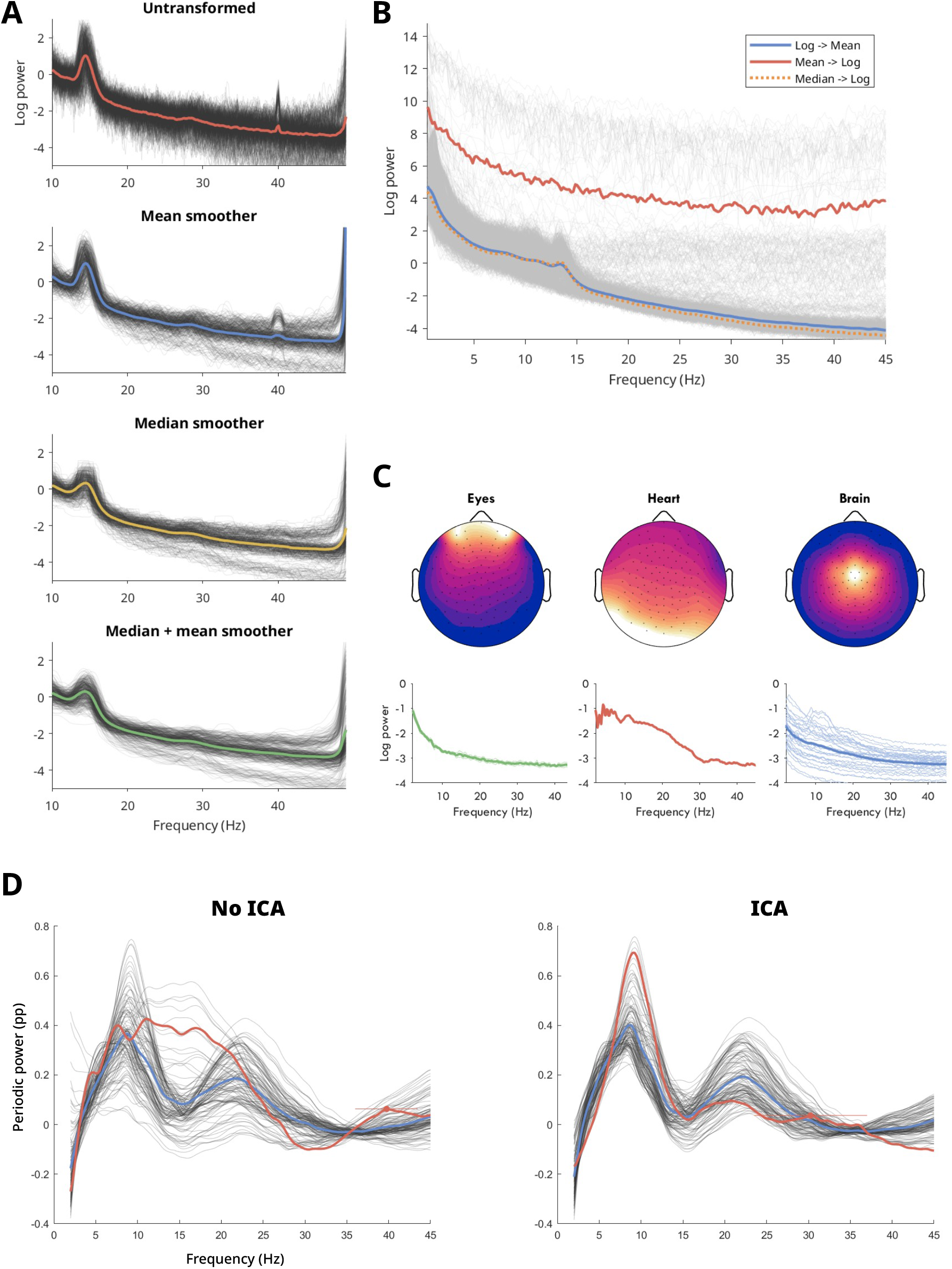
Preprocessing choices. A) Smoothing the spectral signal prior to specparam speeds up processing time and reduces the detection of spurious peaks. However, when a mean smoother is applied, if there are sharp peaks due to environmental electrical noise, these will create broader bandwidth peaks, now indistinguishable from physiological spectral peaks. Applying a median smoother of 2-3 Hz suppresses these narrow electrical spikes while leaving the broader physiological peaks largely intact (albeit at lower amplitudes). However, median smoothers are no longer “smooth” but rather blocky, which again increases specparam computation time. Applying first a median and then a mean smoother to single segment power spectra combines the advantages of both median and mean smoothers. The disadvantage is that the amplitudes of the peaks are reduced. Normally, this smoothing process is not needed for specparam when calculating power with Welch’s method, because power is calculated over longer periods, which means more discrete Fast Fourier Transforms (FFTs) get averaged together and thus producing a smoother signal anyway. B) Artefacts are often several orders of magnitude larger than EEG data, however during sleep they usually only occur occasionally, i.e. during body movements. Log-transforming spectra before averaging them across epochs and/or channels reduces the impact of these artefactual outliers substantially. A similar effect can be obtained by the median (dotted line). C) Examples of components detected using independent component analysis (EEGLAB’s function *pop_runica*). Unlike iota activity, ocular artefacts are in the frontal-most channels, peaking over channels 25 and 8. Heartbeat artefacts are located primarily in edge-occipital left channels. The three spectra below each topography provide the spectra of all ICA components for one wake EEG recording. D) Periodic power during REM sleep with and without ICA; the most substantial change is in the sigma-beta range, due to heartbeat artefacts. These could largely be ignored by removing the outer edge channels.

**Suppl. Figure 4.12:**
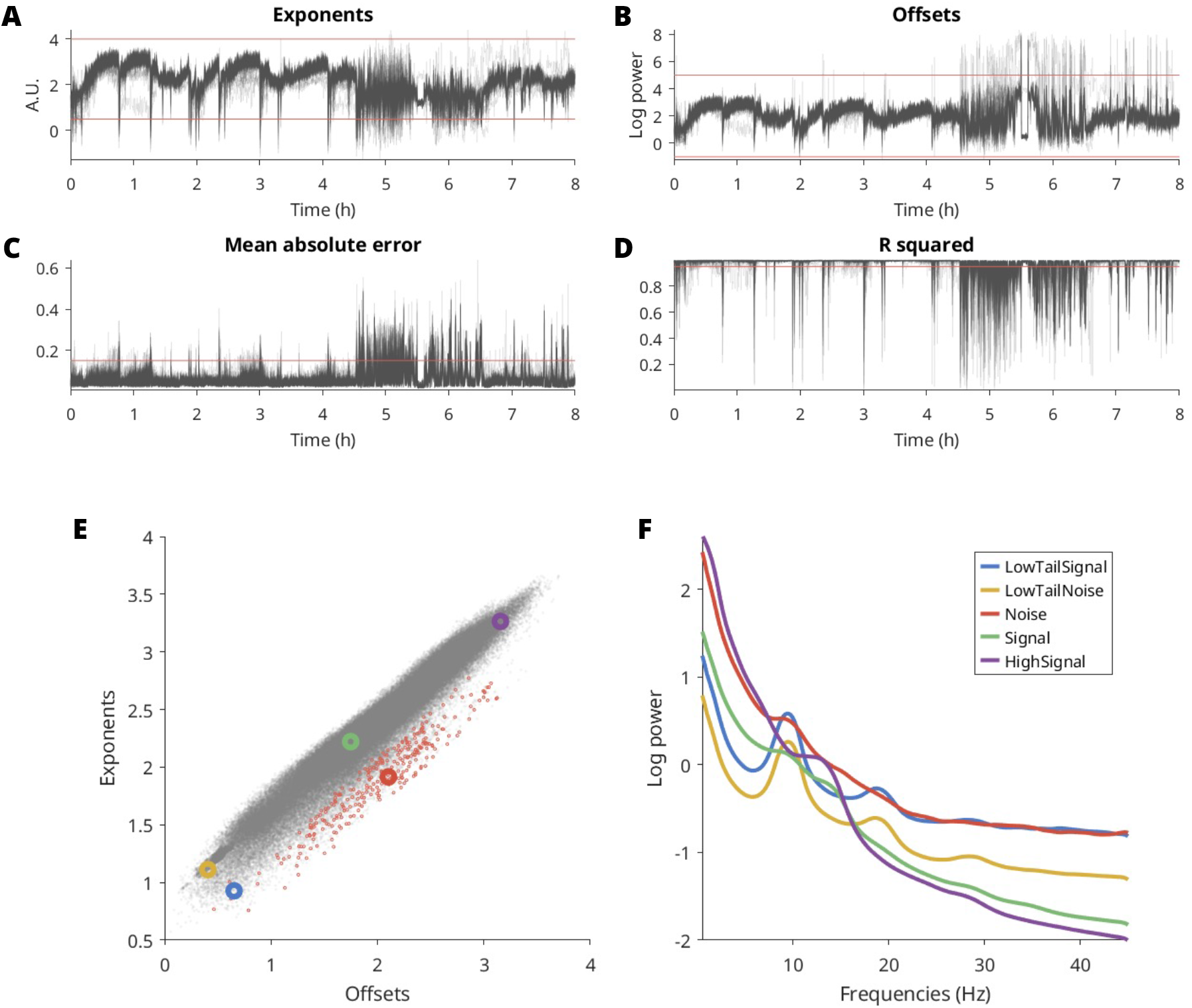
Specparam artefact rejection from a single night. A-D) Basic thresholds on specparam outputs (red lines). A) Aperiodic exponents. Unfortunately, the naturally large fluctuations in exponents necessitate wide thresholds that often do not capture artefactual deviations. Thus, in a second step we removed exponents that deviated from the median by > 0.6 (e.g. see around the 1h mark). B) Aperiodic offsets. C) Mean absolute error (MAE). D) R squared. Note, thresholds are often (but not always) redundant. E) Relationship between exponents and offsets in the example night. Each grey dot is one segment. Red small dots are segments marked as artefacts based on exponent-offset combination. Large circles indicate the mean of a quadrant of data used for F. F) Average log spectra for example quadrants in E. Note how the red Noise spectrum has the same high-amplitude low frequencies as the purple High Signal (mostly N3 segments, as identifiable by the sigma periodic peak), but the same high frequencies as the blue Low Tail Signal (mostly wake, as identifiable by the 10 Hz peak). The yellow circle highlights segments that are a combination of signal and artefact (an unplugged signal, when all channels correlate with each other). In clean sleep data, offsets and exponents are linearly correlated with each other, because when exponents change, the aperiodic signal rotates around a pivot point approximately in the alpha-sigma band (Bódizs et al., 2021), thus systematically affecting the offset. Artefact segments can have this pivot frequency in much lower bands, leading to exceptionally shallow exponents for high offsets, or vice-versa. Therefore, this artefact detector: 1) performs a robust linear correlation (*robustfit*) between offsets and exponents; 2) calculates the residuals between measured exponents and the expected ones; 3) z-scores the residuals (with median and median absolute deviation); 4) marks as artefact any segment that has abs(z) > 10. This process is done first separately for each channel, then on the remaining segments all together. This is done in two steps because the linear relationship between exponent and offset slightly varies across channels, and a clear outlier for one channel may be perfectly normal for the group. The rather relaxed threshold of z > 10 accounts for the fact that wake EEG tends to deviate more from this otherwise highly linear relationship between exponents and offsets without strictly being artefactual. This process may work better if applied separately to each sleep stage, however as much as possible we preferred approaches that did not rely on sleep scoring for preprocessing.

**Suppl. Figure 5.1:**
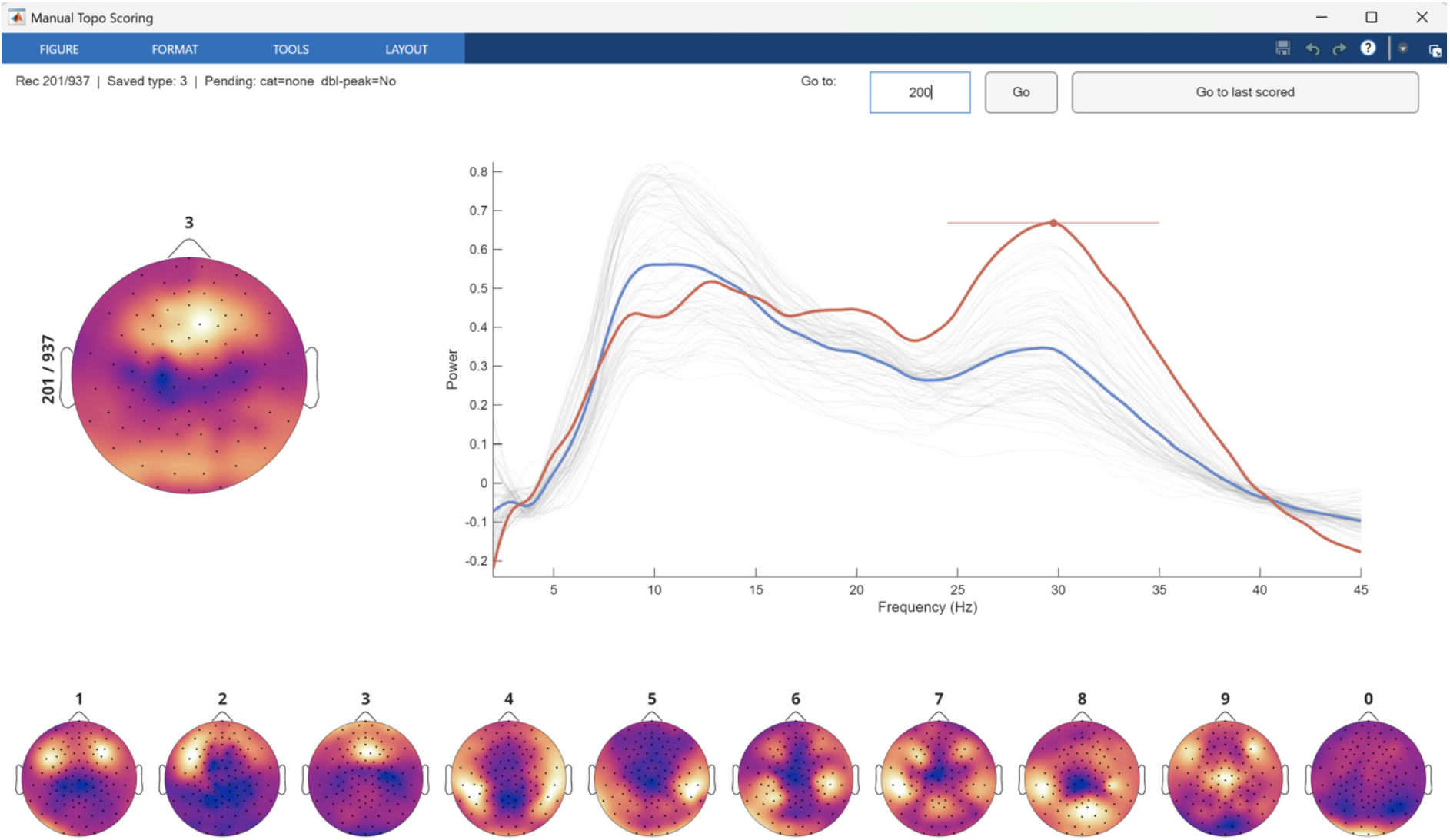
Graphic user interface for scoring iota topographies. Top left topography is the current plot to score, using keys from 0-9, with an example of each provided below. Top right is the spectra with the detected iota peak. User toggles “d” if they see a distinct second (or more) peak.

**Suppl. Table 1:** REM sleep vs not-REM sleep log-power using paired t-tests, from one recording per participant. FDR correction for multiple comparisons was conducted over all comparisons.

| Band | N3 | N2 | N1 | W |
| --- | --- | --- | --- | --- |
| **Theta** | t = -73.40; p = 4.37e-290; d = -1.36; N = 567 | t = -36.51; p = 2.99e-151; d = -0.64; N = 575 | t = 0.06; p = 0.955; d = 0.00; N = 331 | t = 6.76; p = 9.81e-11; d = 0.27; N = 256 |
| **Alpha** | t = -36.52; p = 1.66e-150; d = -1.12; N = 567 | t = -45.24; p = 1.36e-190; d = -1.02; N = 575 | t = -12.38; p = 4.14e-29; d = -0.38; N = 331 | t = -25.83; p = 6.45e-73; d = -1.23; N = 256 |
| **Sigma** | t = -28.32; p = 3.75e-110; d = -0.91; N = 567 | t = -52.17; p = 5.33e-219; d = -1.39; N = 575 | t = -21.70; p = 2.49e-65; d = -0.54; N = 331 | t = -19.69; p = 5.6e-53; d = -0.74; N = 256 |
| **Beta** | t = 40.16; p = 3.25e-167; d = 1.10; N = 567 | t = 20.22; p = 6.95e-69; d = 0.36; N = 575 | t = -13.73; p = 3.64e-34; d = -0.28; N = 331 | t = -32.30; p = 8.31e-92; d = -1.02; N = 256 |
| **Iota** | t = 43.00; p = 9.12e-180; d = 1.06; N = 567 | t = 25.34; p = 2.63e-95; d = 0.43; N = 575 | t = -9.97; p = 1.33e-20; d = -0.18; N = 331 | t = -29.05; p = 1.35e-82; d = -0.72; N = 256 |

**Suppl. Table 2:** REM sleep vs not-REM sleep periodic power.

| Band | N3 | N2 | N1 | W |
| --- | --- | --- | --- | --- |
| **Theta** | t = 1.72; p = 0.091; d = 0.08; N = 567 | t = 7.49; p = 3.95e-13; d = 0.24; N = 575 | t = 1.95; p = 0.0581; d = 0.07; N = 331 | t = -6.61; p = 3.06e-10; d = -0.39; N = 256 |
| **Alpha** | t = -4.98; p = 1.02e-06; d = -0.22; N = 567 | t = -21.69; p = 3.31e-76; d = -0.71; N = 575 | t = -13.12; p = 1.02e-31; d = -0.41; N = 331 | t = -33.16; p = 9.13e-94; d = -1.87; N = 256 |
| **Sigma** | t = -14.26; p = 2.22e-39; d = -0.67; N = 567 | t = -40.89; p = 3.53e-171; d = -1.65; N = 575 | t = -20.78; p = 1.34e-61; d = -0.57; N = 331 | t = -22.35; p = 1.19e-61; d = -0.98; N = 256 |
| **Beta** | t = 51.94; p = 3.53e-216; d = 2.22; N = 567 | t = 29.32; p = 2.42e-115; d = 0.79; N = 575 | t = -5.39; p = 1.65e-07; d = -0.15; N = 331 | t = -25.70; p = 2.52e-72; d = -1.00; N = 256 |
| **Iota** | t = 22.49; p = 5.75e-80; d = 1.20; N = 567 | t = 24.44; p = 2.3e-90; d = 0.86; N = 575 | t = 6.58; p = 2.62e-10; d = 0.18; N = 331 | t = 0.03; p = 0.98; d = 0.00; N = 256 |

**Suppl. Table 3:**
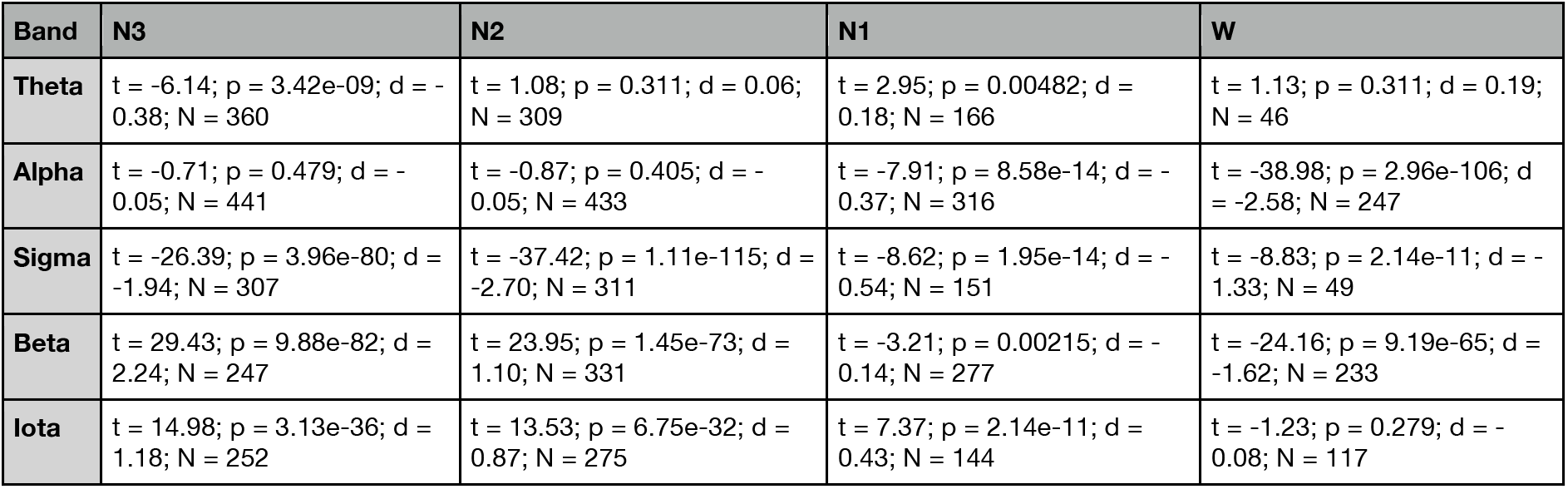
REM sleep vs not-REM sleep periodic peak amplitudes.

